# Inhibiting toxicity of ALS-linked dipeptide repeat proteins by polystyrene sulfonate (PSS)

**DOI:** 10.1101/2023.05.19.541518

**Authors:** Anna Bratek-Skicki, Junaid Ahmed, Joris Van Lindt, Karl Jonckheere, Eveline Peeters, Kara Heeren, Fatemeh Fakhri, Alex Volkov, Jelle Hendrix, Piotr Batys, Peter Nagy, Richard G. Pestell, Ludo Van Den Bosch, Peter Tompa

## Abstract

The GGGGCC (G4C2) expansion in the noncoding region of C9orf72 is the most common genetic cause of amyotrophic lateral sclerosis (ALS) and frontotemporal dementia (FTD). The repeat region is translated into five different dipeptide repeat proteins (DPRs), of which the arginine-rich DPRs (R-DPRs) poly- GR (GRn) and poly-PR (PRn) are highly neurotoxic. In this study, we characterized the protective effect against R- DPR toxicity of polystyrene sulfonate (PSS), an FDA-approved drug applied in hyperkalemia, in biochemical, cellular and iPSC-derived motor neuron (MN) models. We found that PSS, in a length-dependent manner, interacts very tightly with R-DPRs, and releases their bound RNA in R-DPR - RNA mixtures. PSS significantly influences the liquid- liquid phase separation (LLPS) of R-DPRs elicited by RNA and reduces their ensuing cell toxicity in Neuro-2a cells. PSS is cell penetrable, and it can effectively rescue the inhibitory effect R-DPRs on axonal mitochondrial transport in iPSC-derived MNs. Shorter (n < 340) variants of PSS are not toxic either to cells or to mice upon intracerebroventricular injection up to 1 mM concentration. Our results suggest that its polymeric nature endows PSS with an advantageous effect in C9-ALS providing a potential therapeutic tool against this debilitating neurodegenerative disease.

## Introduction

Amyotrophic lateral sclerosis (ALS) and frontotemporal dementia (FTD) are severe neurodegenerative diseases that show significant genetic, pathological and clinical overlap. Both diseases encompass a progressive loss of neurons that usually leads to death within 3 to 5 years after diagnosis [1–3].

About 90% of ALS cases are sporadic (sALS, but even in this group twin studies indicate genetic predisposition), while approximately 10% are inherited (familial, fALS). The most prominent genes affected in fALS are superoxide dismutase 1 (*SOD1*), TAR DNA-binding protein (*TARDBP*, encoding TDP-43), fused in sarcoma (FUS) and C9orf72 [4]. Among these, hexanucleotide (G4C2) repeat expansions (HREs) in the 5’ noncoding region of the C9orf72 gene is the most common genetic cause (40%) of fALS, hereafter referred to as C9-ALS [5]; C9orf72 HREs are also implicated in about 10% of cases.

The HREs may theoretically lead to the disease through three, non-mutually exclusive pathological mechanisms. The loss of function of C9orf72 protein may disturb autophagy pathways and endosomal trafficking [6]. Toxic gain-of-function can occur due to the production of mRNAs containing an aberrant, expanded repeat region [7–9]. However, recent studies emphasize the toxic gain-of-function caused by aberrant, repeat-associated, non- AUG (RAN) translation of C9orf72 mRNA into five dipeptide repeat proteins (DPRs) [10, 11] being the most prevalent disease mechanism. Of the five different DPRs, arginine-containing poly-PR and poly-GR (R-DPRs) have been reported to be highly toxic [9, 12, 13] due to disrupting cellular processes such as nucleocytoplasmic transport, RNA processing and the formation of cytoplasmic and nuclear ribonucleoprotein particles (condensates), such as stress granules (SGs) and nucleoli [14–17]. As they carry a very high net charge, R-DPRs can engage in tight interactions with RNA, they can undergo liquid-liquid phase separation (LLPS) and induce this process in combination with RNA and numerous proteins involved in RNA- and stress-granule metabolism, such as nucleophosmin 1 (NPM1) [18] and Ras GTPase-activating protein-binding protein 1 (G3BP1) [19]. R-DPRs are cell penetrable [14], they might serve as biomarkers of fALS [20], and due to their excessive toxicity, they are primary drug targets, although owing to their highly non-natural, disordered structure, they are considered mostly undruggable. Whereas the gain-of-function mechanisms are also supported by direct observation of G4C2-HRE RNA- and DPR- containing inclusions in patients with C9-ALS [21, 22], the three distinct mechanisms do not have to be mutually exclusive and probably act in concert in driving the disease [23].

There are only a few drugs approved by the U.S. Food and Drug Administration (FDA) to treat ALS, such as Radicava, Rilutek, Tigluik and Nuedexta, all of which work by indirect mechanisms. The active compound of Radicava is edaravone, a drug used to treat stroke [24]. The compound is an antioxidant, and its use rests on the hypothesis that oxidative stress contributes to neurodegeneration [25]. Rilutek and Tigluik incorporate riluzole, a sodium-channel inhibitor and antiglutamatergic compound that delays the onset of ventilator-dependence in some patients, extending life expectancy by a few months [26]. Nuedexta contains two compounds, quinidine and dextromethorphan that can treat pseudobulbar effect (PBA) of uncontrolled episodes of laughing or crying in some patients with ALS and other neurological diseases, but its exact mechanism is unknown [27]. Most recently, an antisense oligonucleotide (ASO) against SOD1 has been approved by FDA/EMA as Tofersen [28–31].

Unfortunately, except for Tofersen and Riluzole, the approved drugs only mitigate symptoms and do not target basic mechanism(s) of the disease. Based on the extreme toxicity and cell-penetrating nature [32–34] of R-DPRs in many different cell types and model organisms, we decided to take a different approach to tackle ALS caused by C9orf72 hexanucleotide-expansions. Quite recently, an “ultrahigh” (picomolar) affinity interaction between two oppositely charged intrinsically disordered proteins (IDPs), histone H1 and prothymosin-α, was reported [35]. By analogy, we hypothesized that intrinsically disordered positive R-DPRs could be targeted by a highly negatively charged polymeric substance, polystyrene sulfonate (PSS) [36, 37]. PSS is an FDA-approved drug for treating hyperkalemia (cf. Suppl. Table S1), and due to its high negative charge, we hypothesized that it would engage in very tight binding with R- DPRs, counteract their LLPS and alleviate cellular toxicity of R-DPRs in disease. Accordanceingly, we could demonstrate that PSS can bind both poly-PR and poly-GR with a K_d_ of single-digit nMs, it interferes with R-DPR LLPS and releases R-DPR-bound RNA and has a protective effect in cellular models of C9orf72-related ALS.

## Results

### Polystyrene sulfonate binds C9orf72 R-DPRs very tightly

Polystyrene sulfonate (PSS) is a negatively charged polyelectrolyte bearing sulfonate groups (Scheme 1). Due to the very low pKa of the sulfonate group (- 2.1), PSS is fully charged in a wide range of pH and ionic strength conditions. PSS is commonly used in additives [38], membranes [39], ion exchange resins [40], and personal care products [41]. PSS is also used as an FDA-approved drug to treat hyperkalemia in patients with renal insufficiency ([42], cf. Suppl. Table S1) and as an adjuvant during hemodialysis [43]. It was also reported that PSS forms stable complexes with positively charged proteins that can be used in layer-by-layer assemblies to improve biological activity [44, 45].

**Scheme 1.**
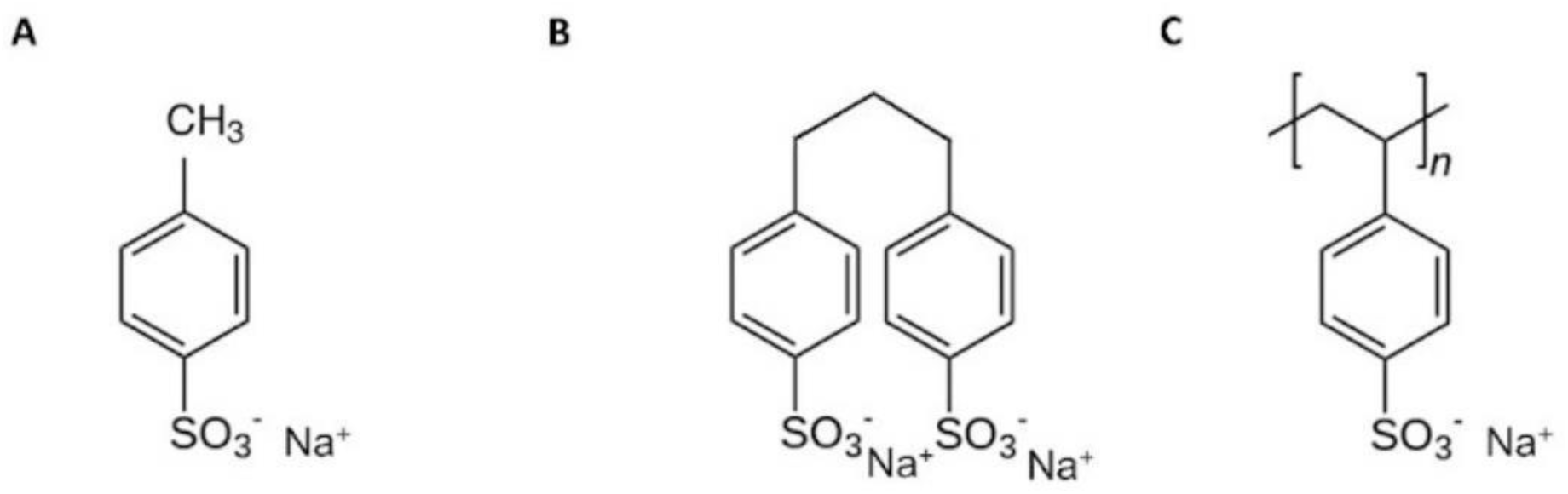
Chemical formula of polystyrene sulfonate (PSS) variants. In this study, variants of polystyrene sulfonate (poly(sodium 4-styrenesulfonate), PSS) of different polymer length were used, such as A, n = 1 (monomer), B, n = 2 (dimer), and longer variants (C), i.e., n = 5 (pentamer), n = 22 (PSS_22) n = 43 (PSS_43), n* = 73 (PSS_Rhod, a heterogenous mixture of average Mw = 15 000 g/mol, fluorescently labelled with Rhodamine), n = 87 (PSS_87) and n = 340 (PSS_340).

In accordance, the very high and opposite net charges of the two polymers (R-DPRs are positive, whereas PSS is negative) suggest a strong interaction between them. In fact, in a chemically similar arrangement, a picomolar Kd between two oppositely charged IDPs, histone H1 and prothymosin-α, was already observed [35]. To address the Kd of the PSS - R-DPR interaction, we studied a whole range of PSS length variants from a monomer (PSS_1) to a 340-mer (PSS_340), as outlined in Scheme 1. To determine the Kd of PSS - R-DPR binding, we first studied a long PSS (PSS_43) and applied isothermal titration calorimetry (ITC), which yielded very steep binding curves with both PR30 and GR30 (Fig. 1A, B), whereas PA30 which has no charge and is of limited toxicity in ALS models [46], gives no detectable heat signal (Fig. 1C). Analysis of PR30 and GR30 curves indicates very strong binding: fitting the data (Fig. 1D, E) yielded apparent Kd values of 12.6 nM and 2.4 nM for PR30 and GR30, respectively, but indicated no detectable binding of PA30 (Fig. 1F).

**Figure 1.**
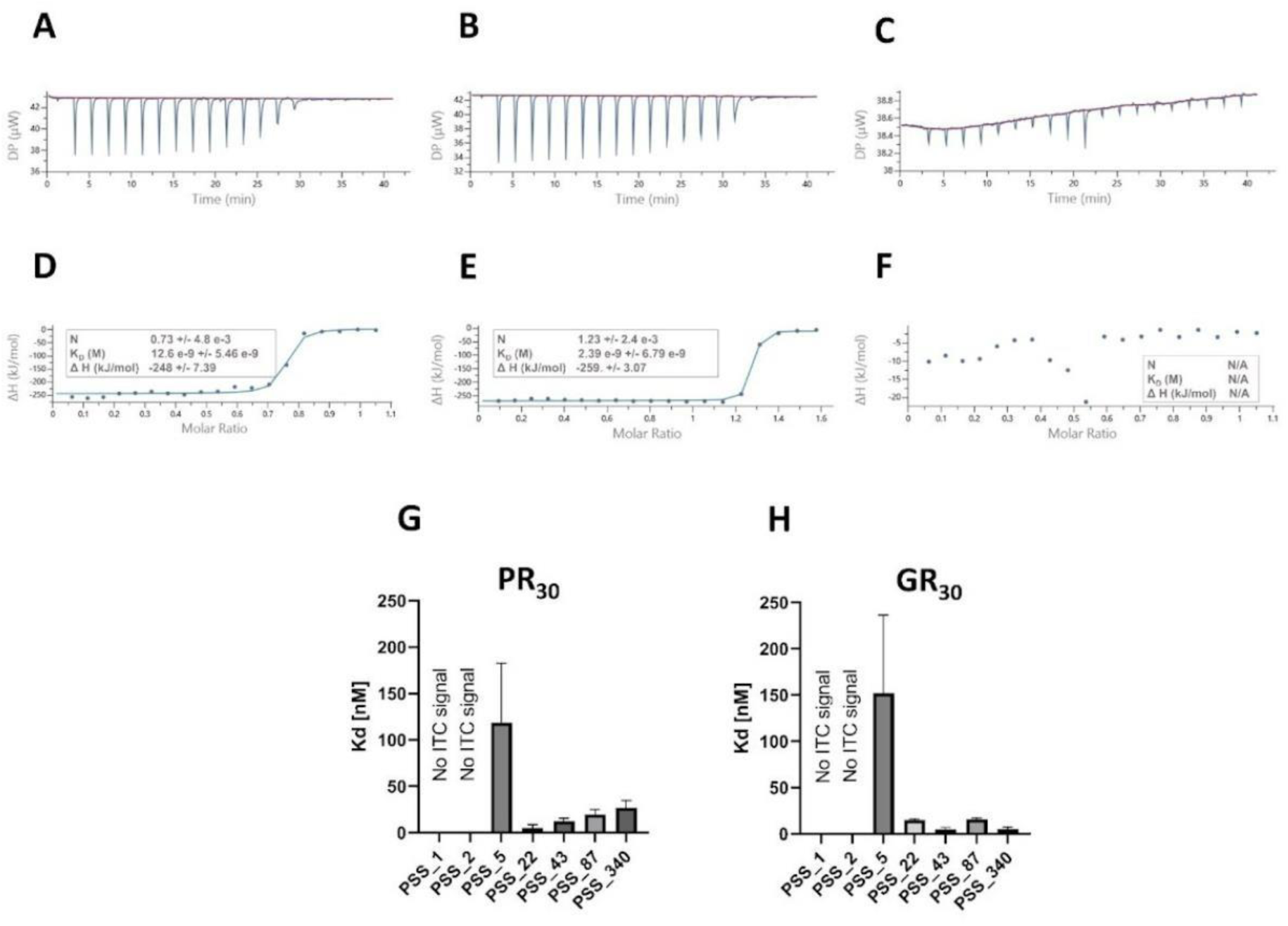
Tight binding of DPRs by PSS. The binding of PSS to DPRs (PR30, GR30 and PA30) was demonstrated and quantified by isothermal titration calorimetry (ITC). Three DPRs were titrated by different length variants of PSS. (A-C) raw ITC data with PSS_43, (D-F) integrated ITC data, with curves fit to the standard single binding site model. (G, H) As PA30 shows no signs of interaction, Kd is determined for PR30, and GR30 with all PSS variants (cf. Scheme 1) in 150 mM NaCl, pH 7.4.

It is to be noted that numerous repetitions of this experiment yielded a binding constant in the range of nM to pM, and to obtain a valid binding constant in ITC, a “c factor” (c = n[M]Kd, where n is the binding stoichiometry, [M] the macromolecule concentration, and Kd the binding constant) must be between 10 and 100 [47]. Values of c that are too low (<10) can sometimes be used to fit Kd but cannot be used to accurately determine stoichiometry (n). Values of c > 1000 can be used to accurately determine n, but not Kd. Since binding stoichiometry in the PSS:R-DPR system is approximately 1 and PSS_43 concentration in the ITC cell is 20 µM, the c factor is 252,000 (Fig. 1D) and 48,000 (Fig. 1E), i.e., the accurate determination of Kd is probably beyond the limits of ITC. A further reduction of protein and ligand concentrations is not feasible since the signal-to-noise ratio becomes too low.

Therefore, to assess further the very strong interaction between PSS_43 and the R-DPRs, we implemented microscale thermophoresis (MST) measurements, in which fluorescently labelled PR30 and GR30 were titrated with PSS_43 (Suppl. Fig. S1). Here also, very steep binding curves were obtained for both DPRs. The obtained data analysed using the Hill-equation and resulted in strong binding constants 5.7 nM and 6.0 nM for PR30 and GR30, respectively.

We also applied fluorescence correlation spectroscopy (FCS) to analyze the binding of fluorescently labeled DPRs to PSS_43 and compared the deduced binding constants to those obtained using other techniques (ITC, MST) (Suppl. Fig. S2). In these experiments Kds were in the range of nanomolar; 5.2, 7.3, 9.3 nM based on rotational diffusion, fluorescence anisotropy and fluorescence lifetime.

Given the very strong interaction with PSS_43, we next investigated how the Kd changes with polymer length with PR30 (Fig. 1G) and GR30 (Fig. 1H). In both cases, the monomer and dimer show no measurable interaction, PSS_5 already binds with a Kd = 120 nM (PR30) and 150 nM (GR30), whereas longer variants bind significantly more tightly. PSS_22 has the full interaction strength, which varies a little with increasing length (Table 1, Fig. 1G, H). Given the expectation that PSS – R-DPR interaction is primarily driven by electrostatic attraction, we also carried out a salt titration of the PR30 – PSS_43 and GR30 – PSS_43 (Suppl. Fig. S3) system. As expected, the interaction is strongest at 0 mM NaCl and becomes gradually weaker as salt concentration increases from 150 mM to 500 mM.

### PSS inhibits the phase separation of R-DPRs

Stress granules (SGs) involved in pathological protein aggregation in ALS contain proteins with domains that can undergo and drive LLPS. The most notable of these, RNA-binding domains (RNA recognition motifs, RRMs), and intrinsically disordered arginine-rich sequences (RGG boxes) that can also bind RNA or poly(ADP-ribose), initiate LLPS and lead eventually to the aggregation of prion-like domains [48–50]. It has been suggested that arginine-rich motifs can play a significant role in LLPS in both physiological and pathological processes [49, 50]. C9orf72-encoded R-DPRs have been linked with C9-ALS etiology [46] and suggested to exert their pathological effects primarily by impairing the formation of physiological nucleoli [18] and SGs [19]. In recent studies [17, 18, 51], it was suggested that R-DPR LLPS induced by crowding and RNA are reasonable approximations of the complex pathological processes. Therefore, we hypothesized that the LLPS system provides a reasonable approach for the initial assessment of the potential favorable effects of PSS variants in pathological situations.

Having established the very strong binding of PSS to R-DPRs (Fig. 1), we next investigated whether PSS inhibits the RNA-promoted LLPS of R-DPRs. We first initiated LLPS of the studied DPRs by four different RNA variants: polyU, polyA, tRNA and total RNA extracted from Neuro-2a cells (Fig. 2). By following LLPS by turbidity (OD 340 nm), the signal of PR30 (Fig. 2A) and GR30 (Fig. 2B), but not PA30 (Fig. 2C) first increased rapidly, then slowly decreased, probably due to maturation (ripening) of droplets. This behaviour was typical for polyU and polyA, whereas relatively smaller turbidity values were observed for total RNA, suggesting the formation of bigger droplets. A different kinetic trajectory was recorded in the presence of tRNA, where, after a rapid increase, a slow decrease and rather constant OD value was observed. This behavior was consistent with earlier observations that LLPS is promoted by disordered (polyU, polyA) but not ordered (tRNA) RNA molecules [53].

**Figure 2.**
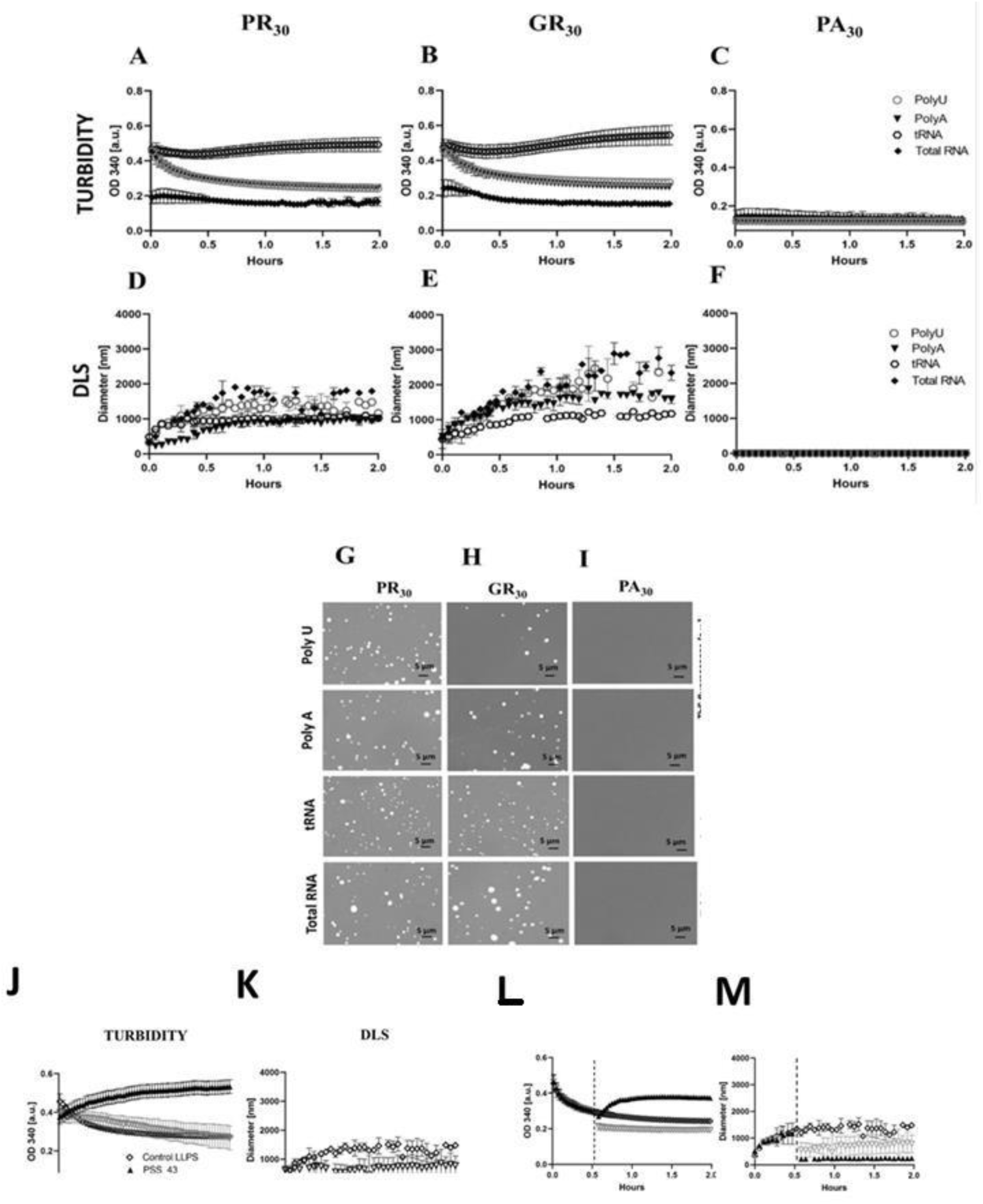
Liquid-liquid phase separation of C9orf72 DPRs initiated by different RNAs. (A-F) LLPS of three different DPRs, PR30 (left column), GR30 (middle column) and PA30 (right column) was monitored by an increase in turbidity (OD340, A to C), and size of droplets by dynamic light scattering (DLS, D to F) in the presence of polyU, polyA, tRNA and total RNA extracted from Neuro-2a cells. Droplets were also visualized by fluorescence microscopy using DyLight 488-labelled DPRs mixed into 200x excess of unlabeled DPRs, following 15 min incubation (PR30 (G), GR30 (H), PA30 (I)). The experiments were performed in 10 mM HEPES, pH 7.4, 150 mM NaCl, at 50 µM DPR and 0.5 µg/µl RNA concentrations (K-N). LLPS of PR30 was then initiated by polyU in the presence of PSS_43 or the monomer of PSS, and monitored by adding PSS before (K, L) or 30 min after (M, N), initiating LLPS, by turbidity (OD340, on K, M) or DLS (on L, N). The experiments were performed at 150 mM NaCl, pH 7.4, at 50 µM PR30 and 0.5 µg/µl RNA concentration. The molar ratio of PSS_43 and the monomeric version was 1:43, which corresponds to the number of styrene sulfonate units in PSS_43.

We then studied the evolution of droplets by dynamic light scattering (DLS). For PR30 and GR30 (Fig. 2D, E) the droplets grew slowly, reaching a maximum in size (approximately 1-2 µm) in about 1 h. In agreement with turbidity measurements, smaller droplets formed in the presence of tRNA for PR30 and GR30, whereas bigger droplets appeared in the presence of total RNA. Some difference was apparent between the two R-DPRs, as GR30 droplets showed slower growth than those formed of PR30. PA30, on the other hand, (Fig. 2F) showed no DLS signal, confirming the turbidity data proving the lack of LLPS in its case.

These conclusions were also confirmed by direct visualization of droplets by fluorescence microscopy (Figure 2G, H and I). Solutions of PR30 and GR30, but not of PA30, quickly formed many droplets upon the addition of all studied RNAs. Typically, the biggest ones appeared with total RNA and the smallest ones with tRNA, in accordanceance with turbidity and DLS measurements. Overall, the results with the three different techniques are in line with the idea that R-DPRs undergo LLPS in the presence of different RNAs, following somewhat different kinetic trajectories.

In pathological conditions, liquid droplets proceed toward aggregated states, as indicated by the fluorescent indicators thioflavin T (ThT) or thioflavin S (ThS, in the presence of RNA [52, 53]). To address whether the various R-DPR - RNA systems show this behavior and thus can be considered as reasonable proxies to pathological LLPS mimicking C9-ALS processes, we next followed the ThS fluorescence signal over 15 h in different R-DPR – RNAs systems (Suppl. Fig. S4). PR30 and GR30, similarly to TAR DNA-binding protein 43 (TDP-43), a protein strongly implicated in the formation of pathological inclusions in ALS [54], developed intensive ThS signals with all RNA variants, polyU, polyA, tRNA, and total RNA (Suppl. Fig. S4A, B). PA30, which does not promote LLPS of R-DPRs, and is only moderately toxic in ALS models, did not show an increase in ThS fluorescence over an incubation time of 15 h.

Given the strong length dependence of PSS – R-DPR binding (Fig. 1, Table 1) and anticipating that the polymeric nature of PSS plays an important role in its effect on R-DPR LLPS, we first compared the influence of PSS_43 and sodium p-toluenesulfonate (monomer of PSS, cf. Scheme 1) on LLPS at identical monomer concentrations (i.e., applying PSS_1 at 43x concentration to that of PSS_43). We initiated LLPS of PR30 under these conditions by polyU and added PSS variants either before the initiation of LLPS or 30 min after polyU addition, following LLPS by turbidity (OD 340) (Fig. 2 K, M) and DLS (Fig. 2L, N) measurements, also confirmed by microscopy (inserts). In all cases, strong interference of LLPS by polymeric PSS was observed. When PSS_43 (at a molar ratio of 1:1) was added to PR30 solution before polyU, this first resulted in a rapid initial jump, followed by a slow rise in absorbance (Fig. 2K), which is opposite to the classical LLPS kinetic trajectory observed in the absence of PSS. DLS (Fig. 2L) showed the immediate formation of small condensates (around 200 nm, as opposed to 1-2 µm without PSS), which were stable over time. In bright-field microscopy (where 200 nm particles are too small to resolve), a small population of larger particles of about 1 µm in diameter was observed (Fig. 3, inserts).

**Figure 3.**
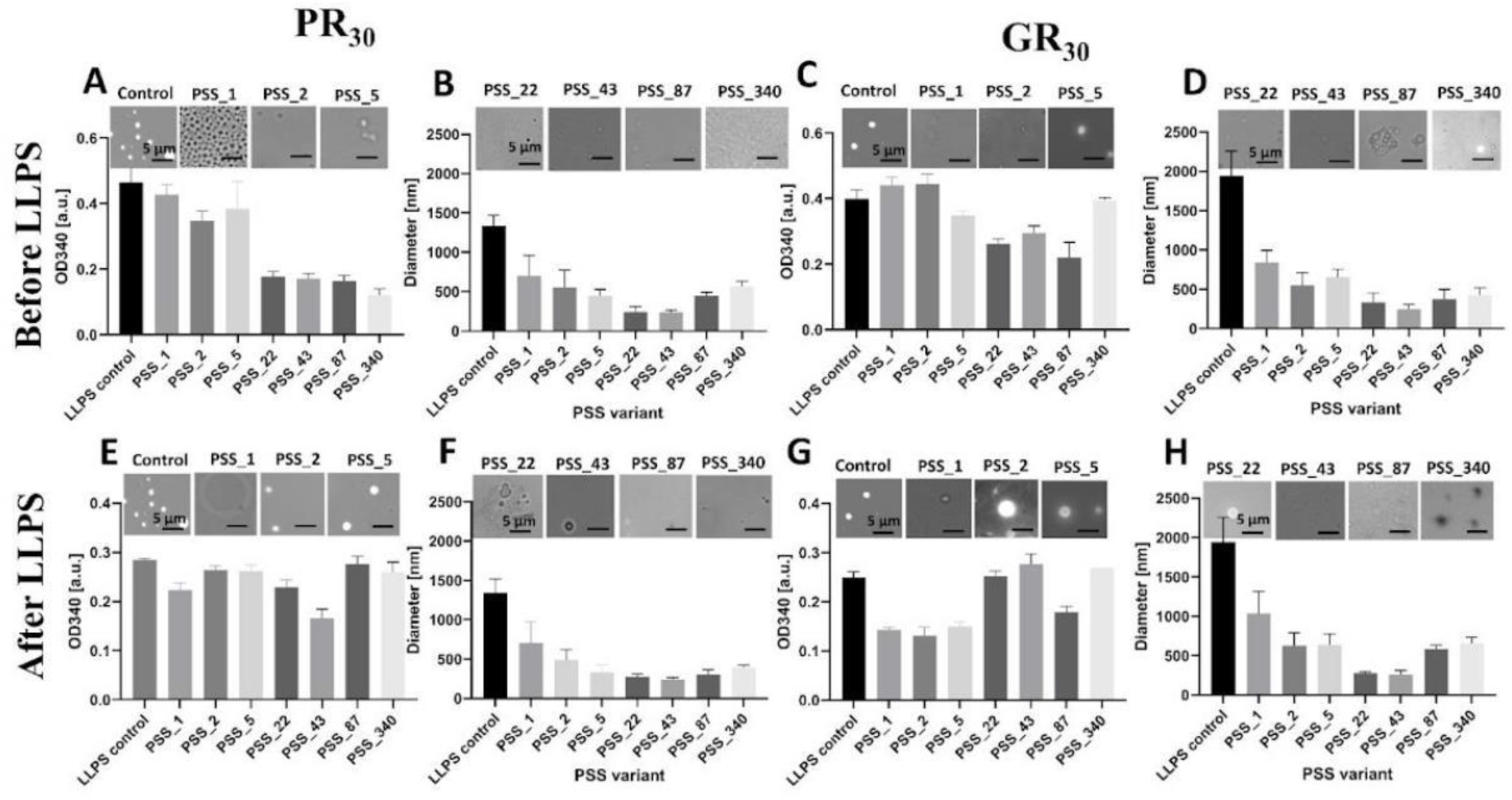
Length-dependence of inhibition of the LLPS of R-DPRs by PSS. LLPS of PR30 (A B, E, F) and GR30 (C, D, G, H) was initiated by polyU in the presence of different length variants of PSS, and monitored by turbidity (OD 340) and DLS, and visualized by microscopy (inserts), by adding PSS before (A-D) or 30 min after (E-H) initiating LLPS. The experiments were performed at 150 mM NaCl, pH 7.4, at 50 µM DPR and 0.5 µg/µl RNA concentration and at a PSS unit to DPR ratio 1:1.

Monomeric PSS_1 applied at 43x the concentration of PSS_43 (Scheme 1A) had a basically different, although not negligible, effect, The increase and decrease in turbidity were slower than in the control experiment run without PSS. In accordance, DLS measurements showed a significantly slower formation of droplets of approximately 800 nm in diameter (Fig. 2L), in between the size of droplets formed with and without PSS_43. This observation was fully supported by microscopy images (Fig. 3, inserts) Moreover, on a different focusing plane, another population of big liquid-like structures was present, not detectable by DLS.

A similar set of experiments was performed for PR30 solutions in which PSS_43 or PSS_1 was added 30 min after the induction of LLPS (Fig. 2M, N). The polymer had an immediate, profound effect. It reversed the direction of turbidity change, resulting in a slow increase in turbidity to a constant value of about 0.4 (Fig. 2M). DLS measurements revealed an immediate cessation of the growth of droplets and their dispersion to small ones of about 200 nm in diameter (Fig. 2N). Microscopy images show only a few small droplets whose size was either too high, or their concentration was too low to be detected by DLS (Fig. 3, inserts).

When the monomer PSS_1 was added to an already phase-separating solution of PR30, it hardly affected the turbidity signal. DLS showed a drop in droplet size, to around 800-1000 nm, confirming an effect that is much less than that of PSS_43. Microscopy also supported this conclusion: droplets of µm size are observed together with big, apparently liquid structures detectable in a different focusing plane.

As very similar behavior was observed for the GR30 solution (Suppl. Fig. S5). Therefore, we conclude that polymeric PSS has a dramatic effect on the LLPS propensity of R- DPRs. In this effect, the polymeric nature of PSS is of utmost importance, as the application of the same number of units in the form of a monomer has much less effect.

### Polymeric nature of PSS is important for its effect on R-DPRs

Given the importance of polymer length in R-DPR binding (Fig. 1) and in potential applications against R-DPR toxicity, we next scanned the whole range of PSS variants (cf. Scheme 1) for their inhibitory effect of R-DPR LLPS. In these experiments (Fig. 3), LLPS of PR30 or GR30 was initiated by polyU, and PSS length variants (always at concentrations to ensure a PSS unit to DPR ratio of 1:1) were added before or after the initiation of LLPS (adding polyU) and LLPS was quantitated by the maximal value of its OD340 turbidity curve (as on Fig. 2A and B). When the polymer was added before LLPS, the critical length for inhibition is between n = 5 and 22 with both R-DPRs (Fig. 3A, C). When the PSS is added after LLPS, there is no apparent effect (Fig. 3E, G), but it must be considered that turbidity is not a good indicator of LLPS inhibition (cf. Fig. 2). When measured by DLS, even a PSS monomer had a significant effect in all scenarios (whether with PR30 or GR30, added before or after polyU), and the full effect is reached at n = 22 (reducing droplets to about 1/5 of their original size), above which increasing polymer length had no further effect (Fig. 3B, D, F, H). Microscopic images support this conclusion (Fig. 3, inserts), although they cannot be as well quantified and are complicated by the occasional appearance of much larger condensates. In conclusion, the length n = 22 is close to the length of DPRs (n = 30), which may be of importance in reaching a maximal effect, serving as a guide in our further search for PSS variant(s) most effective in countering R-DPR toxicity.

### PSS releases tightly bound RNA from R-DPRs

To get mechanistic insight into the observed strong effect of PSS on the RNA-induced LLPS of R-DPRs elicited by their very strong binding, we carried out electrophoretic mobility shift assays (EMSA) experiments with the R-DPR – RNA – PSS system. In particular, we investigated whether RNA bound by R-DPRs can be released by PSS due to a tighter PSS – R-DPR than RNA – R-DPR interaction. First, we determined the Kd of the RNA–R-DPR interaction, we titrated 32P-labelled mRNA with either PR30 (Fig. 4A, B) or GR30 (Fig. 4C, D). Therefore, we determined the amount of unbound and bound RNA by densitometry following autoradiography of the unbound (lower apparent Mw) and R-DPR-bound (higher apparent Mw) RNA bands. Fitting the titration curves by a 1:1 binding model, we obtained the Kd for RNA-PR30 (125 nM) and RNA- GR30 (70 nM) interaction. Apparently, they are almost two orders of magnitude weaker than the PSS binding to R-DPRs.

**Figure 4.**
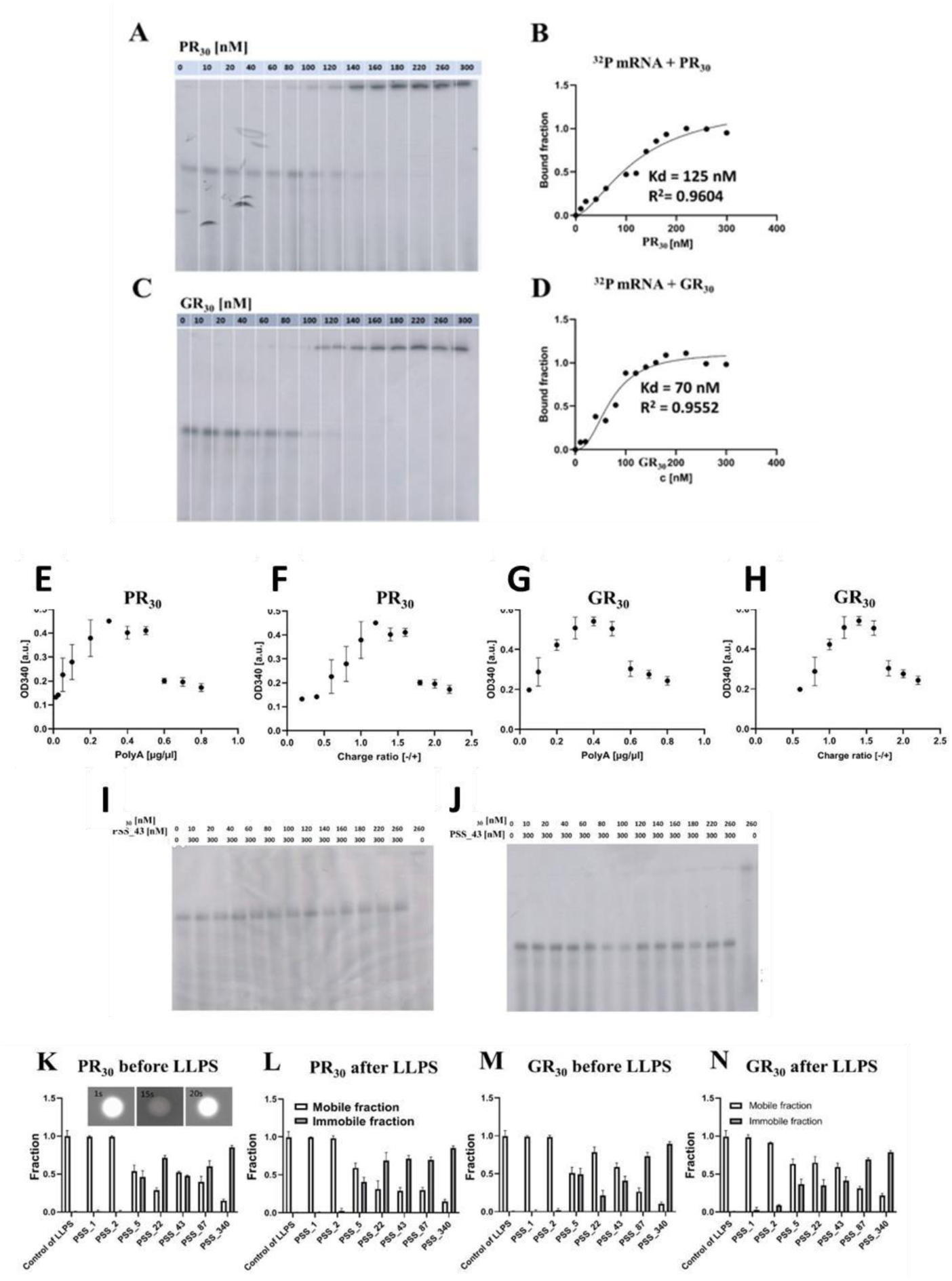
Mechanism of LLPS and PSS effect on DPRs. EMSA results show strong interactions between 32PmRNA and PR30 (A, B) and GR30 (C,D). The figures show the fraction of 32PmRNA band as PR30 (A, B) and GR30 (C, D) were titrated. The values for the Kd and Bmax obtained from the curve fitting are 125.4 +/- 1.2 nM and Bmax 1.307 +/- 0.05 for PR30 and 70.11 +/- 8.2 and Bmax 1.118 +/- 0.08 forGR30. RDPs titrated by polyA. The maximum of recorded turbidity (OD340) curves plotted as a function of polyA concentration (E, G) or RNA:R-DPR charge ratio (F, H) shows a maximum, i.e., reentrant behavior. Upon titrating 32PmRNA in the presence of an excess of PSS_43 (300 nM), the RNA always appears in the free (non-DPR complexed) state, except when no PSS is added (I, J). R-DPRs were treated with different variants of PSSs added before (K, M) or 30 min after (L, N) LLPS was initiated and % FRAP recovery (the fraction of fluorescence recovered, i.e., dynamic fraction of LLSP droplets), was determined; empty columns show fast recovering, whereas full columns show non-recovering, fraction of fluorescence.

As LLPS of R-DPRs is promoted by RNA [17], we next investigated how the amplitude of LLPS, as measured by peak turbidity (cf. Fig. 1), changes upon titration of R-DPRs with an excess of RNA. In the case of both PR30 (Fig. 4E) and GR30 (Fig. 4G), LLPS shows a reentrant behavior [55, 56], increasing initially but becoming inhibited at the excess of RNA, which is in line with a model assuming optimal LLPS at stoichiometric amounts of the two oppositely charged components. In fact, when LLPS intensity curves are plotted as a function of the charge ratio of RNA versus DPRs, the maximum of LLPS is achieved in both cases (Fig. 4F, H) at 1:1 charge ratio, above which LLPS becomes inhibited by excess RNA (and dominance of its negative charge).

RNA drives LLPS of R-DPRs probably by establishing crosslinks between R-DPR – RNA clusters [17], as also suggested for the LLPS of G3BP1 [57]. In this scenario, the tighter binding of PSS to R-DPRs might suggest that the inhibition of RNA-driven LLPS b y PS S stems from releasing RNA from R-DPRs. Therefore, we next investigated whether PSS added to the RNA - R-DPR mixture can liberate RNA from the complex. As RNA binds tightly to R-DPRs, resulting in its apparently very slow release from the complex and equilibration with PSS, we first mixed PSS_43 and RNA at equimolar concentrations, and then added R-DPRs PR30 (Fig. 4I) and GR30 (Fig. 4J) in increasing concentrations. RNA appeared in isolation every time PSS_43 was in molar excess to the R-DPR, with the RNA-R-DPR complex appearing on the gel only in the absence of PSS.

These results support the model that the inhibitory effect of PSS on RNA - R-DPR LLPS emerges from PSS terminating RNA binding, while itself not being able to promote a PSS - R-DPR phase separation. This is not evident, as R-DPR – PSS droplets could form similarly to R-DPR – RNA droplets, due to similar charge complementarity in the two systems. Whereas DLS and microscopy experiments (Fig. 2) suggest that this is not the case, we have further approached this issue by fluorescence recovery after photobleaching (FRAP) experiments, which give direct information on the dynamics of condensates. In fact, when measuring the FRAP recovery of R-DPR - polyU droplets, they appear fully mobile, underscoring their liquid nature compatible with a model of formation by LLPS (Fig. 4K - N). When PSS of different length is added, whether before or after initiating LLPS (causing the appearance of much smaller droplets, cf. Fig. 2 and Fig. 3), an interesting trend appears: at lengths PSS_1 to PSS_2, which have an overall smaller effect, the droplets are fully mobile, suggesting little interference by PSS. At PSS_5, and then fully from PSS_22 onwards droplets become very rigid, their fraction of recovery dropping to about 30% or less. This means that longer PSS variants not only inhibit LLPS and affect droplet morphology to a great extent, they also basically reduce the dynamics of droplets (condensates) that replace liquid droplets.

### Molecular mechanisms of R-DPR – PSS interaction

Our results suggest not only that PSS can engage in much stronger interactions with R-DPRs than RNA, but also that in an atomistic-structural sense these interactions critically differ so that not even the very long PSS_340 (Fig. 5K-N) can support the formation of liquid droplets. To understand this mode of interaction in atomistic detail, we carried out experiments and modelling to characterize the R- DPR – PSS interaction. We applied NMR spectroscopy to characterize which regions of the polymers engage in contacts upon interaction. As both R-DPRs and PSS are highly repetitive, causing severe signal overlap in multi-dimensional NMR, we recorded 1D 1H NMR spectra of PSS_43 and PR30 (Fig. 5A, B) and GR30 and PA30 (Suppl. Fig. S6) at increasing R-DPR concentration. The spectra of all three DPRs were then also acquired in the presence of increasing concentrations of PSS_43 (Fig. 5C, D).

**Figure 5.**
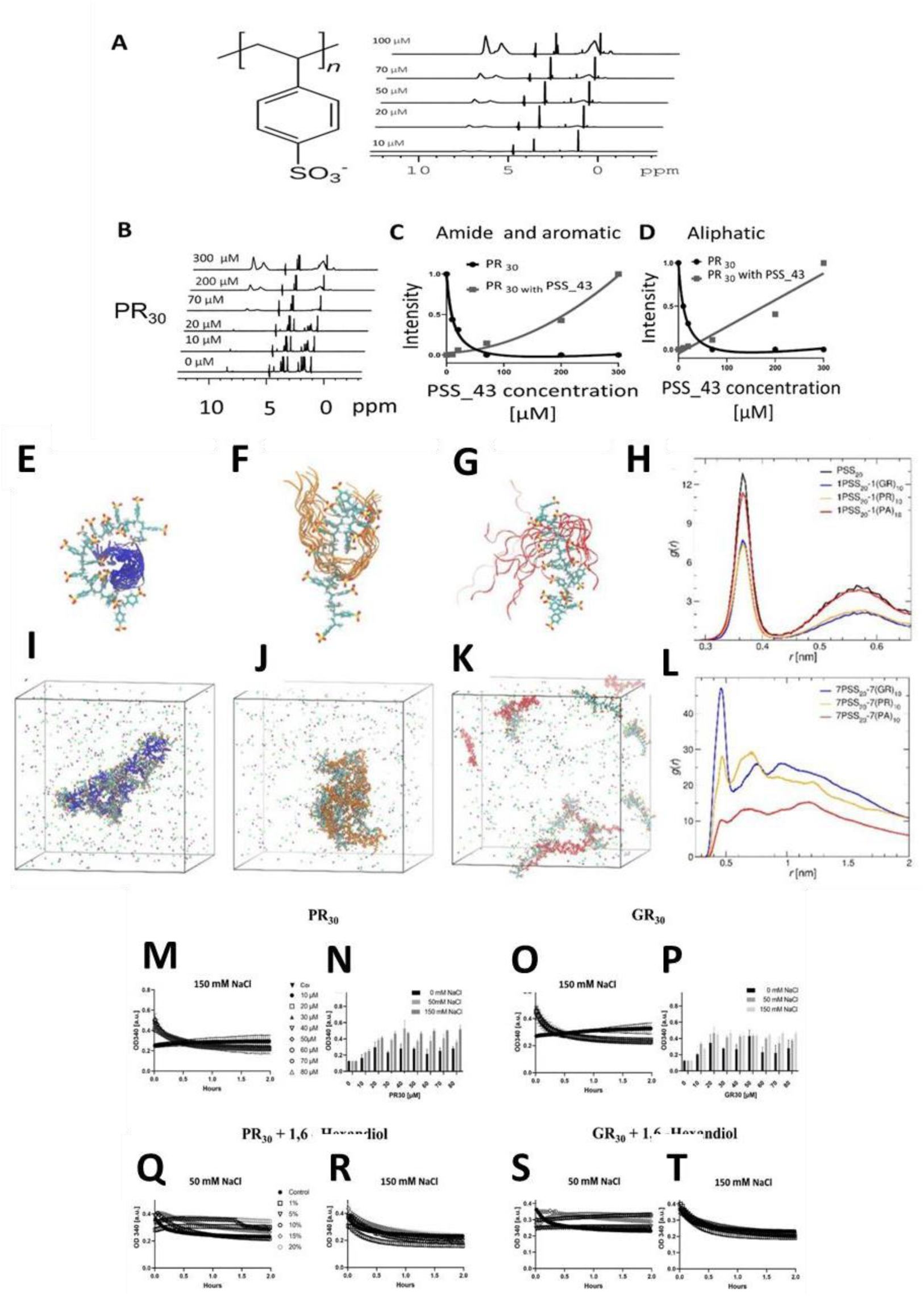
Mechanism of PSS - R-DPR interaction. ^1^D^1^H NMR Spectrum of PSS_43 (A) was recorded at different concentrations; peaks correspond to the chemical structure of PSS. The interaction of PSS with DPR PR30 (B - D) was followed by ^1^D-^1^H NMR: spectra of PR30 at 100 μM were recorded in the presence of increasing PSS_43 concentrations (B, up to 300 μM). Peak intensities in the aromatic and amide (6-9 ppm) and aliphatic (1-3 ppm) regions were integrated and plotted separately for the DPR and PSS as a function of PSS concentration (C, D). MD simulation snapshots of 1 PSS_20 : 1 GR10 (E), 1 PSS_20 : 1 PR10 (F) and 1 PSS_20 : 1 PA10 (G) systems, aligned to the PSS_20 molecules. Peptide conformations, saved in 20 ns time intervals, are shown as ribbons. The radial distribution function (RDF) between the sulphur atoms of PSS and Na+ counterions in the mixtures and a single PSS_20 molecule as a reference, is shown (H). Snapshots after 300 ns of 7 PSS_20 : 7 GR10 (I), 7 PSS_20 : 7 PR10 (J) and 7 PSS_20: 7 PA10 (K) systems. The Na+ and Cl- ions are highlighted as green and pink spheres, respectively. The RDF between the Cα atoms of DPRs and the sulphur atoms of PSS are calculated (L); RDFs are averaged over three different initial configurations. All simulations were run at 150 mM NaCl concentration. LLPS of PR30, and GR30 were monitored by turbidity (OD340) at different concentrations of DPRs (M-P), three ionic strength values (I = 0, 150, 500 mM) or five concentrations of hexanediol (Q-T). The concentration of DPR for the experiments with hexanediol was 50 µM, RNA in all experiments was added at 0.5 µg/µl.

The spectrum of PR30 has a peak in the amide region at 8.4 ppm. The addition of 10 µM PSS_43 results in a decrease in intensity of this peak, whereas at 70 µM PSS_43 the amide peak completely disappears and the intensity of the PSS_43 peak starts to increase. The peaks of PSS appear at 7.4 and 6.6 ppm, corresponding to the aromatic ring of PSS [58]. In the aliphatic region, multiple peaks for PR30 resulting from spin- spin coupling can be observed. Upon the addition of PSS, a similar binding behavior was observed as in the amide region, the intensity of the backbone of PSS_43 [58] starts to appear at about 70 µM PSS. The interaction of PSS_43 and PR30 is based on a slow exchange of protons; this data was fitted with 1:1 binding, because tight binding fitting did not result in reliable dissociation constant (Fig. 5B-D).

In the case of GR30, two peaks appear in the amide region, at 8.6 and 8.35 ppm. Their intensity decreases upon the addition of 10 µM PSS_43, and the peaks completely disappear at 100 µM PSS_43. The data were fitted by 1:1 binding, because slow- exchange fitting did not result in a reliable dissociation constant (Suppl. Fig. S6AE-C).

In the case of PA30, a peak in the amide region was observed at 8.2 ppm, which decreases upon the addition of 20 µM PSS_43, upon further addition of PSS_43, it starts to decrease. We can conclude that the interaction is weak and is based on a fast exchange, with multiple binding events involved in the interaction between PSS and PA30 (Suppl. Fig. S6D-F).

Overall, the interaction of PSS_43 and R-DPRs is based on a slow exchange, whereas the interaction with PA30 is based on a fast exchange that involves multiple binding events. With PR30 and GR30, both the backbone (amide region) and side chains (aliphatic region) are involved in binding.

As NMR cannot provide high-resolution structural information for the R-DPR – PS interaction, we carried out all-atom molecular dynamics (MD) simulations to characterize the system further. Single molecules, i.e., PSS_20, GR10, PR10, and PA10 (for feasibility, DPRs shorter than n = 30 were modeled) were equilibrated and analyzed. The obtained conformations and secondary structure evolutions are presented in Suppl. Figs. S7-S10. These configurations were used further to construct multiple-molecule systems, i.e., containing PSS and peptide pairs (1:1 system) and more complex mixtures containing 7 copies of each molecule (7:7 system). The MD behavior of the multi-molecule systems is outlined in Fig. 5E-L, and Suppl. Figs. S11-S19, and Suppl. Tables S2 and S3.

The molecular dynamics results highlight significant differences between R-DPRs GR10 (Fig. 5E) and PR10 (Fig. 5F), showing strong association with PSS20, and PA10 (Fig. 5G) that engages only in very weak/transient interactions with the charged polymer. The interaction between these peptides and PSS leads to the release of counterions condensed around the PSS molecule, which is manifested in decreasing peak height of radial distribution function (RDF) between the S atoms and Na+ ions (Fig. 5H): such an effect is suggestive of a primarily electrostatic component driving poly- PR – PSS and poly-GR – PSS interactions, as already suggested for polyelectrolyte complex formation [61], and supported by the salt dependence of their PSS_43 binding (Suppl. Fig. S3). PSS also forms a hydrogen-bond network with GR10 and PR10 (Suppl. Tables S2 and S3), which is more extensive than the one with PA10.

MD simulations of multicomponent (7:7) systems (Figs. 5I-L) confirm these predictions of atomic contacts observed in the 1:1 systems and underscore the different LLPS tendencies of R-DPRs (cf. Fig. 2). That is, both GR10 (Fig. 5I) and PR10 (Fig. 5J), but not PA10 (Fig. 5K), form dense clusters with PSS, quantitatively confirmed by RDF values between Cα atoms of DPRs and sulfur atoms of PSS (Fig. 5L). This behavior is in line with the observed formation of small residual clusters from R-DPR – RNA droplets upon the addition of PSS (Fig. 2), as also underscored by their reduced dynamics observed by FRAP (Fig. 4K-N).

MD simulations can further assess the electrostatic component of interactions, as simulations at different salt concentrations can be carried out and compared. A strong salt effect upon varying NaCl concentration appears (Figs. 5E-L show results of 150 mM NaCl concentration; results of higher salt are shown in Suppl. Fig. S11). RDFs indicate that the release of counterions condensed around PSS is significantly weaker at higher NaCl concentrations, with a similar trend observable for hydrogen bonding (Suppl. Table S2) and the number of contacts between the peptides and PSS (Suppl. Fig. S19), all these trends underscore the primarily electrostatic nature of interactions.

To provide further experimental confirmation of these inferences, we have carried out salt titrations of R-DPR – PSS interaction by ITC and salt and 1,6-hexanediol titration of R-DPR – RNA LLPS by turbidimetry. In both systems, salt has a strong inhibitory effect on LLPS of both PR30 and GR30, even at physiological, 150 mM concentration. Interestingly, 1,6-hexanediol, an agent known to interfere with hydrophobic interactions, is hardly effective at 150 mM NaCl, suggesting that electrostatics is the dominant force in R-DPR LLPS, hydrophobicity playing a secondary role.

### PSS is of limited cell toxicity and can penetrate cells

Motivated by the very effective inhibition of the LLPS of R-DPRs by PSS and the suggested effect of R-DPRs on biomolecular condensation in the cell [19], we wanted to move toward cellular applications of the polymer, in particular to testing if it can avert R-DPR-induced toxicity. Therefore, we first asked if PSS itself is toxic to cells. While PSS is taken at very high doses in hyperkalemia, like 60g/day [36, 37], it might pass through the digestive track without being absorbed.

First, we incubated Neuro-2a cells with all different PSS length variants (from n = 1 to 340) for 24h and assessed cell viability by total ATP content (Fig. 6A). We found that with the exception of the longest, PSS_340 (IC50 = 5.3 ± 6.8 μM), none of the other PSS variants decreased viability. In comparison, we have also incubated Neuro-2a cells with the three dipeptide repeats (Fig. 6B), and found that GR30 (IC50 = 4.9 ± 2.4 μM) and PR30 (IC50 = 12.8 ± 7.7 μM) are highly toxic, whereas PA30 has a very moderate effect on cells (IC50 > 30 μM), in accordance with similar experiments carried out on different other cell lines [59, 60]. Whereas R-DPRs are known to be cell penetrable [18, 61], it has never been tested if PSS could penetrate the cell membrane to engage with R-DPRs within cells. Therefore, we tested two types of cells by fluorescent microscopy, adding Rhodamine-labelled PSS_73*_Rhod (a heterogeneous mixture), PSS_5_Rhod and PSS_ 24_Rhod (cf. Scheme 1) to Neuro-2a cells (Fig. 6C), and PSS_73*_Rhod and PSS_43_Rhod to U2OS cells (Fig. 6D). Images taken after 24 h incubation show intracellular localization of all PSS variants in both cells. When we compared the localization of PSS_73*_Rhod and DyLight 488-labelled GR30 or PR30 inside cells (Fig. 6E), we observed the colocalization of R-DPR and PSS molecules, as expected by their very tight binding. 4,6-Diamidino2-phenylindole (DAPI) staining for DNA showed a preferential cytoplasmic localization of both PSS and DPR molecules. It was already shown that while PR30 is predominantly localized in the nucleus, GR30 is found in both the nucleus and the cytoplasm [64]. As a consequence, we concluded that when PSS is also present, it changes the preferential localization of both R-DPRs to the cytoplasm (Fig. 6E). Their physical association in cells, given the lack of PSS-driven R-DPR LLPS (Fig. 2M), strongly indicates that PSS can counter R-DPR toxicity in cells. We addressed this next.

**Figure 6.**
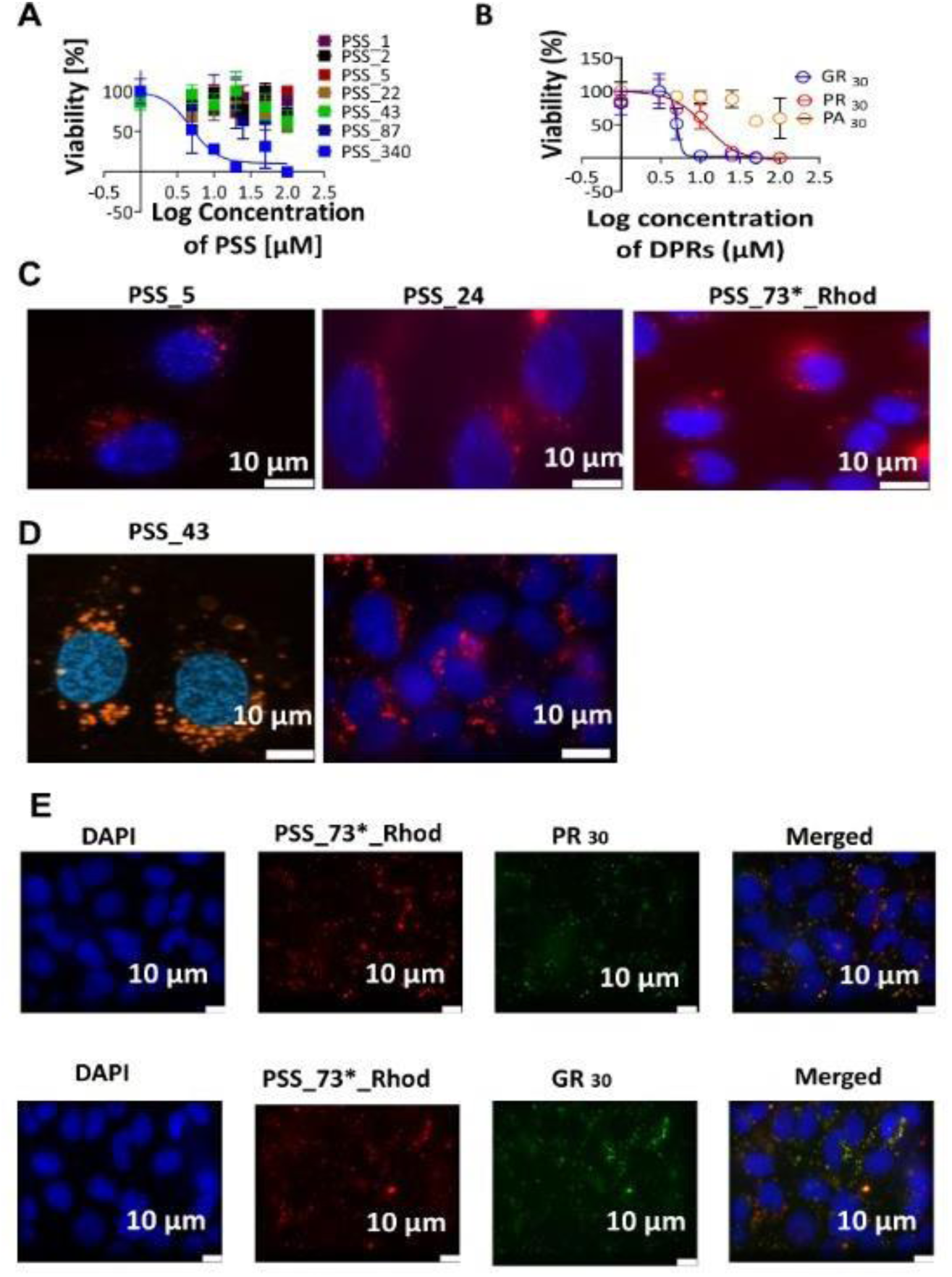
PSS toxicity, cell penetration, and colocalization with R-DPRs. Neuro 2a cells were treated with (A) different PSS length variants (from N = 1 to 340) and (B) different DPRs (PR30, GR30, and PA30) for 24h at concentrations up to 100 μM. Viability of cells following the treatment was assessed by total ATP content, yielding inhibitory concentrations (IC50): 5.3 ± 6.8 μM (PSS_340), 4.9 ± 2.4 μM (GR30), 12.8 ± 7.7 μM (PR30) and >30 μM (PA30). Shorter PSS variants appeared non-toxic under the given conditions. Potential cell penetration of PSS was addressed with fluorescence microscopy by fluorescently labelled PSS_73*_Rhod (a heterogeneous mixture), PSS_5_Rhod and PSS_ 22_Rhod in Neuro 2a cells (C) cells and PSS_73*_Rhod and PSS_43_Rhod in U2OS cells (D), at 10 μM concentration (or 0.12 μg/ml for PSS_73*_Rhod). Images were taken after 24 hr incubation. Colocalization of PSS with DPRs (E) was assessed by incubation of U20S cells with 0.12 μg/ml PSS_73*_Rhod for 2 h followed by 5 μM or 2 μM DyLight 488 NHS-labelled PR30 or GR30, respectively. Images were obtained after 24hr incubation. Cells were also labelled with 4,6-diamidino2-phenylindole (DAPI) for double-stranded DNA.

### PSS is a strong inhibitor of cellular R-DPR toxicity

Because of its very tight R-DPR binding, efficient LLPS inhibition, lack of cell toxicity, cell penetration and cellular colocalization with R-DPRs, we next studied the effect of PSS on cell toxicity of R-DPRs. In accordance with the literature [62], we have shown that both PR30 and GR30 are very toxic to cells measured by total ATP content (Fig. 6B), and we also observed that they severely affect the morphology of Neuro-2a cells and decrease the total number of cells in cell culture (Fig. 7A, B). As suggested by ATP content measurements, GR30 is more toxic than PR30, whereas PA30 is not toxic to cells.

**Figure 7.**
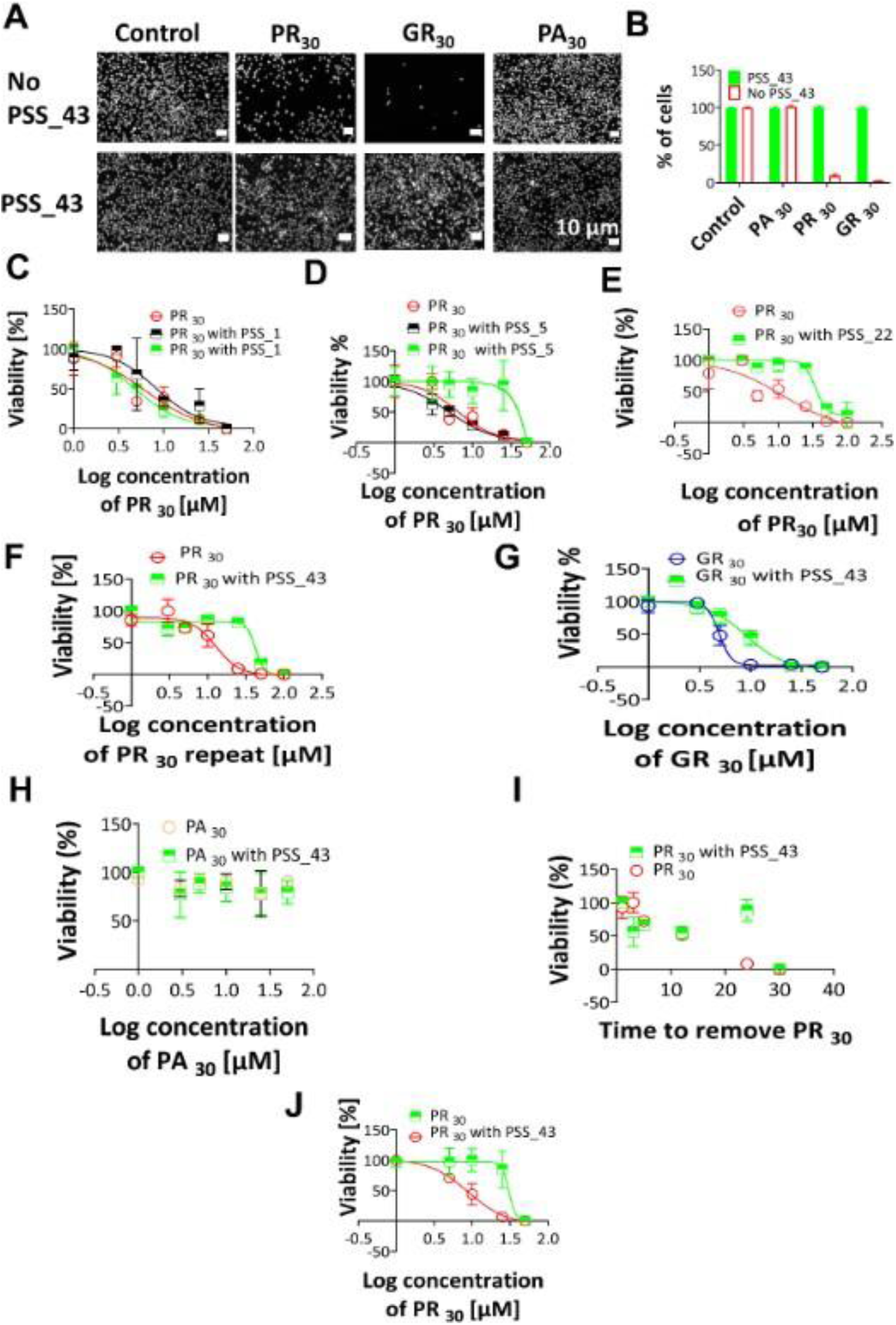
Protection of cells by PSS against R-DPR toxicity. Neuro 2a cells were first incubated without (control) or with three DPRs at 10 μM, in the absence or presence of 10 μM PSS_43. Microscopic images were recorded after 24 h incubation (A), from which total cell counts were determined (B). The protective effect of PSS against the toxicity of both PR30 and GR30 was also measured by adding the DPR at the indicated concentration together with PSS_1, PSS_5, PSS_22, PSS_43 at 20 μM and 200 μM, and ATP content of cells corresponding to viability was measured after 24 h incubation (C, D, E, F, respectively). In a similar experiment (G), PSS_43 at 20 μM concentration was also incubated with GR30 at the concentration indicated and ATP content was measured after 24 h. (H) A similar experiment was also performed with PA30). The protective effect of PSS_43 was also measured by pre-incubating cells with 10 μM PR30 for the time indicated, removing it in a washing step and adding 10 μM PSS_43. Total ATP content was then measured after further 48h of incubation (I). In an experiment of the reverse order, cells were preincubated with 20 μM PSS_43 for 2 h, then PR30 was added at different concentrations indicated, and the ATP content was measured after 24 h incubation (J).

Most importantly, we found that the toxicity of R-DPRs is reversed by PSS. When Neuro 2a cells were incubated with PR30 or GR30 at different concentrations, together with 20 or 200 μM PSS_1, we observed very little cell protection (Fig. 7C). PSS_5, on the other hand, is already very effective at 200 μM concentration (Fig. 7D), whereas upon the application of only 20 μM of the longer PSS_22 and PSS_43, the total ATP content measured after 24 h incubation shows a diminution of R-DPR toxicity, i.e., a significant increase in the effective IC50 values of R-DPRs (to 36.47 ± 6.7 μM (PSS_22) and 42.1 ± 10.4 μM (PSS_43) in the case of PR30, and 9.0 ± 3.1 μM (PSS_43) in the case of GR30 (Fig. 7E, F, G). A similar experiment was performed with PA30, where no effect was observed in the presence of PSS_43 (Fig. 7H).

The very tight binding of PSS to R-DPRs prevents their entry into cells. Therefore, we preincubated cells with 10 μM PR30 (for the times indicated, when R-DPRs enter cells, Fig. 6I). Subsequently, cells were washed to remove the DPR and PSS_43 at 10 μM was added. Following further 48h of incubation, the total ATP content (indicative of viability) of cells (Fig. 7I) showed that cells were affected by toxic PR30, but the ones incubated with PR30 for a longer period (20-30h) were saved by PSS_43, even though PSS and DPR were never in physical contact outside the cell. This observation can be rationalized by two scenarios, either that (i) PR30 can equilibrate through the cell membrane and is effectively dragged out of cells by external PSS, or (ii) PSS penetrates the cell membrane and physically encounters the toxic R-DPR inside cells. To address these possibilities, cells were also preincubated with 10 μM PSS_43 for 2h (when it enters cells cf. Fig. 6), then PR30 at the concentrations indicated was added, and ATP content measured, after 24 h incubation (Fig. 7J). Clearly, cells after long incubations experienced much less toxicity, attesting to the protective effect of PSS.

### PSS rescues R-DPR-induced mitochondrial transport defects

We also wanted to demonstrate that PSS added externally can protect a disease-linked cellular model. Therefore, we have carried out experiments to demonstrate that PSS can rescue R-DPR axonal transport defects, a common motor neuron phenotype associated with ALS. First, we measured the toxicity of PSS_22 on C9or72-derived iPSCs differentiated into MNs. We found that PSS_22 is not toxic at concentrations 5 mM, 10 mM and 25 mM (results not shown), so we used it at 5 mM and 10 mM concentration in the rescue experiments, to be on the safe side.

Mitochondrial transport in live iPSC-derived motor neurons was detected using time-lapse microscopy and three different measures were quantified: (i) average speed of moving mitochondria, (ii) number of moving mitochondria, and (iii) total displacement of mitochondria, as described earlier [63]. Axonal transport defects were induced by incubating the motor neurons with R-DPRs for 8 h, as described previously [64]. Next, to determine whether PSS_22 can rescue transport defects, we removed the R-DPR-containing media and replaced it with non-toxic doses of PSS_22 (0 µM, 5 µM and 10 µM). This experimental approach ensures PSS_22 and GR30 do not interact outside the cells.

As expected, incubating motor neurons with GR30 induced significant mitochondrial transport defects in all measures, which were completely rescued by 10 µM of PSS_22, (Figure 8B). A concentration-dependent rescue could be observed, as average mitochondrial speed defects could already be rescued at 5 µM of PSS_22. Importantly, PSS_22 alone had now effect on mitochondrial movement in non-GR30 treated motor neurons, showing that it specifically rescues GR30-induced mitochondrial transport defects (Figure 8A).

**Figure 8.**
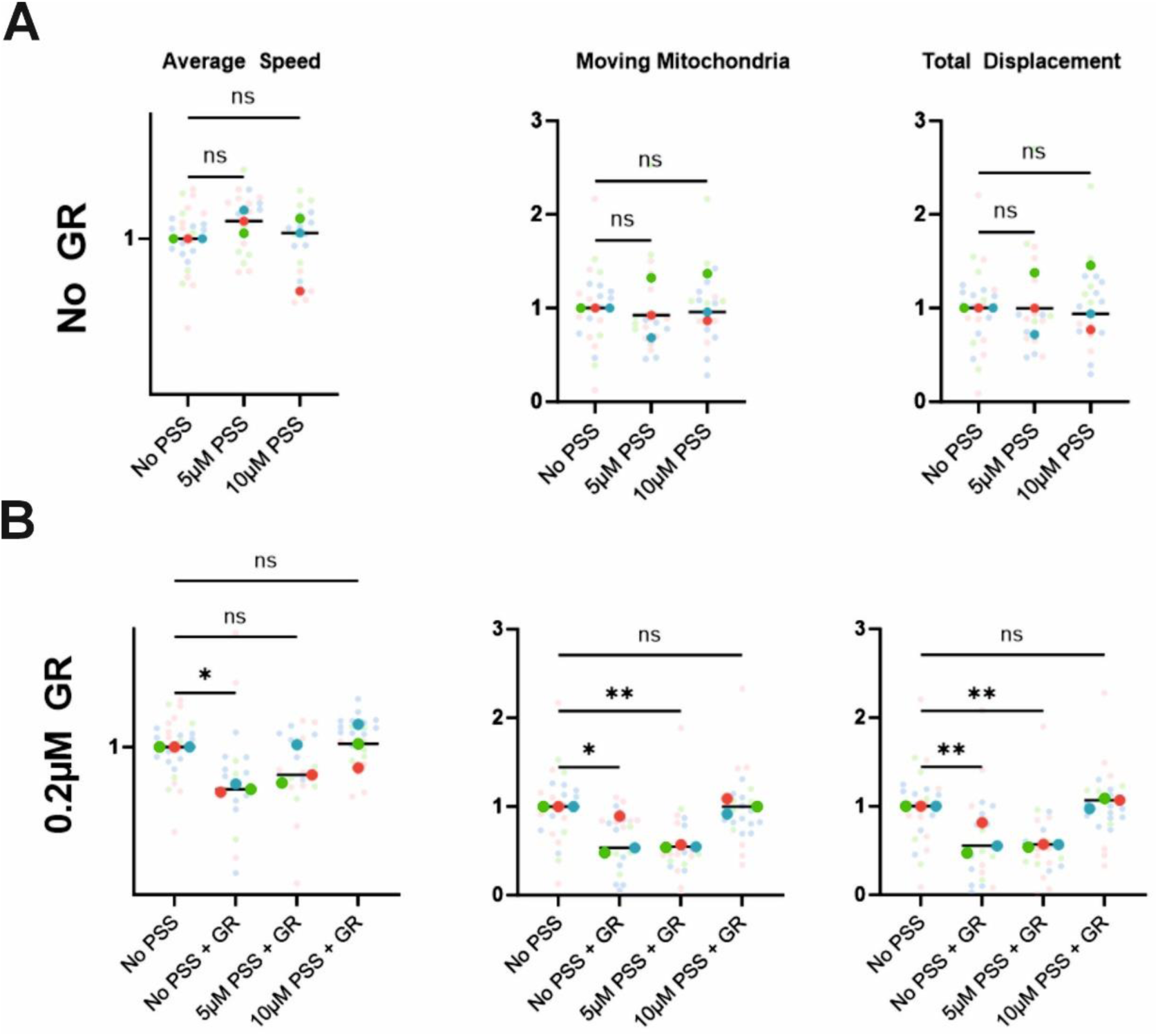
PSS rescues R-DPR-induced mitochondrial transport defects. Mitochondrial transport in iPSC-derived MNs was visualized by time-lapse microscopy [63]. The transport was quantified with three measures: (i) average speed of moving mitochondria, (ii) number of moving mitochondria, and (iii) total displacement of mitochondria moving, on cells not treated (A) or pre-treated (B) with 0.2 mM GR30. Per condition, three biological replicates (each consisting of 5-10 technical replicates) were used to perform statistical analysis. To determine significance, Dunnett’s multiple comparisons test was used. ns = non-significant, * = p<0.05; ** = p<0.01

### Limited mouse toxicity of PSS

Mouse toxicity analysis of PSS_22 upon intracerebroventricular (ICV) injection was assessed at 1 mM and 10 mM concentrations. A toxicity score was given by assessing weight (loss), seizure, atypical behavior, hyperactivity, and normal grooming/eating of 5 mice at each concentration.

The results showed that mice injected with phosphate-buffered saline (PBS) vehicle and 1 mM PSS_22 did not exhibit weight loss during a 7-day follow-up period and their toxicity score did not exceed 1 on a scale of 50, indicating that 1 mM PSS_22 injection is safe for a one-week period. On the other hand, mice injected with 10 mM PSS_22 exhibited signs of toxicity with a mean score of 44.8 directly after the surgery. These results suggest a safe ICV application window of PSS_22 up to 1 mM in mouse, which sets the stage for future protection and efficacy studies.

## Discussion

Neurodegenerative diseases, such as Alzheimer’s disease, Parkinson’s disease or ALS represent an almost unbearable burden on our ageing society, against which we have only very limited medical tools so far. We mostly rely on symptomatic treatment to delay the occurrence of symptoms (most effective in Parkinson’s disease). A special challenge with ALS is that it is a very rapidly progressing disease, leading to death within 2-5 years after diagnosis [4, 65, 66]. A recent, inspiring breakthrough in ALS field is that Tofersen, an ASO against SOD1-ALS [27] has been recently approved by FDA, while an antisense oligonucleotide (ASO) against C9orf72 mRNA has reached clinical trials [29–31]. More help is unquestionably needed, in which PSS may offer a remedy coming from a truly unexpected chemical angle.

C9orf72 R-DPRs are thought to be the primary culprits in neuronal cell death in C9-ALS [18, 54, 67], shown to be highly toxic in cellular [68, 69] and animal [62, 69–71] models of the disease. Their mechanism of action is not fully clear, but impairment of LLPS processes leading to the disruption of cellular RNA metabolism and the dynamics of ribonucleoprotein condensates, such as SGs and nucleoli, could play a critical role [14–16]. Due to their high arginine content, poly-PR and poly-GR can actually penetrate the cell membrane and be toxic to cells when added externally [14, 18].

Due to their central role in C9-ALS, these R-DPRs are potentially very valuable drug targets [29–31]. Because of their intrinsically disordered, extremely non- natural character as proteins, they are considered mostly undruggable. IDPs in general have mostly resisted drug development efforts and are subject to intense scrutiny for rational or phenotypic drug-development efforts [72]. Our results with PSS may represent a novel paradigm in IDP targeting. In every relevant aspect, we have observed its high potentcy, as it can: (i) bind R-DPRs very tightly, (ii) release their bound RNAs, (iii) inhibit R-DPR LLPS, (iv) enter cells on its own, while (v) having limited toxicity. Most importantly, (vi) it curbs R-DPR toxicity in cells.

Although we have made a broad range of relevant observations, the cellular mechanism(s) of PSS protection needs further scrutiny, as it may act in various ways. As demonstrated, the most toxic R-DPRs (poly-PR and poly-GR) are cell penetrable, which may suggest that they can equilibrate across cell membranes, although very tight RNA binding may keep them inside cells. Of course, if an even more potent binder, PSS, is added in the extracellular space, it may alter this equilibrium and cause a net outflow of DPRs from cells, thus reducing their toxic load in the disease. Another possibility is that - as we have shown – PSS can enter cells, co-localize with R-DPRs, reduce their effective intracellular concentration and/or alter their RNA binding and LLPS propensity, while not promoting R-DPR – PSS LLPS on its own.

The observation that PSS can effectively counter LLPS of DPRs, which is critical in the pathogenic mechanism affecting the balanced formation and dispersion of cellular ribonucleoprotein condensates, such as SGs and nucleoli [14–16], and impairing axonal transport [63] in affected cells. We can anticipate that it is central to PSS efficacy that the polymer actually binds R-DPRs much tighter than RNA, thus it can favorably compete with pathogenic mechanisms realized through R-DPR – RNA interactions, and possibly revigorated impaired RNA metabolism, such as splicing, translation, and especially condensation and cellular transport.

A special aspect of the use of PSS is that it is an FDA-approved drug [42, 73], thus its repurposing for C9orf72-related ALS cases may represent an opportunity to develop an effective drug. It was approved in 1958 [73], and has since been used broadly, in a variety of products (Suppl. Table S1). Its successful targeting R-DPRs as a novel modality of countering C9-ALS is validated by recent progress of anti-C9orf72 mRNA ASOs, which decrease R-DPR expression and entered clinical trials [29–31].

It is not to be left without consideration that PSS can be toxic upon particular ways of administration, which may need to be further worked out, as its penetration of the blood-brain barrier (BBB) has never been tested and assessed. Our results pointing to limited toxicity in cells, in iPSC-derived MNs and even in mice, are encouraging for further pre-clinical animal studies. Moreover, it is of specific interest that toxic R-DPRs are involved not only in ALS but also other diseases induced by repeat expansions. To mention a few, poly-PR is involved in spinocerebellar ataxia type 36 (SCA36) [74], poly-GR in X-linked dystonia parkinsonism (XDP) [75], and poly-QAGR in myotonic dystrophy type 2 (DM2) [76], while toxicity of highly positively charged proteins and peptides has been noted in a broad range of infections and diseases, considered as “polycationic poisoning” [77]. The potential application of PSS as a remedy, therefore, may also be explored in other indications.

## Materials and Methods

### Chemical reagents

Sodium p-toluenesulfonate (PSS monomer, cf. Scheme 1), 95%, and poly(sodium 4-styrenesulfonate), PSS Mw = 70 000 (PSS_340) were purchased from Sigma Aldrich while PSS of Mw = 1 000 (PSS_5) and Mw = 8 890 (Mw/Mn = 1.09) (PSS_43) were bought from Polysciences and Scientific Polymer Products Inc., respectively. Rhodamine-labelled PSS (heterogeneous mixture of an average Mw = 15 000) was bought from Surflay Nanotec. HEPES, Tris buffers, NaCl Tween-20, EDTA N,N,N′,N′-Tetramethyl ethylenediamine (TEMED) were purchased from Aldrich. Boric acid and ammonium persulfate (APS) were bought from VWR company while Acrylamide from BioRad.

### Peptides and RNAs

PR30, GR30 and PA30 dipeptide repeats were synthesised by Synpeptide Co., Ltd company, Shanghai, China. Polyuridylic acid (PolyU, cat. no. P9528) potassium salt a heterogeneous and Polyadenylic acid (PolyA, cat. no. P9403) with molar mass ranging between 700-3500 kDa were purchased from Sigma Aldrich. Yeast tRNA (tRNA) was purchased from Thermo Scientific (cat. no. 15401011). Total RNA was extracted from Neuro 2a cells using Trizol from Fisher Scientific, (cat. no.15-596-026) and accompanying protocol.

### Liquid-liquid phase separation (LLPS)

Liquid-liquid phase separation (LLPS) of C9orf72 DPRs was initiated in a buffer of 10 mM HEPES, pH 7.4, at various NaCl concentrations (0 mM, 50 mM and 150 mM). The concentration of DPRs varied from 10 µM to 80 µM, whereas the concentration of RNAs varied from 0.05 to 0.8 µg/µl. LLPS was induced by the addition of RNA and was followed by either turbidity measurements or dynamic light scattering (DLS).

### Turbidity measurements

LLPS was followed by monitoring the change of the turbidity of the solution in a BioTek SynergyTM Mx plate reader in which non-binding black 96 well plates with transparent bottoms (Sigma®) were used. Solutions were mixed, the plate covered with a transparent film (VIEWsealTM), and the absorbance of the solution was monitored at 340 nm for 2 hours at 25°C with continuous shaking. The experiments were conducted in triplicate and then averaged.

### Dynamic light scattering (DLS)

Dynamic light scattering (DLS) measurements were carried out on a DynaPro NanoStar (Wyatt) instrument. A disposable cuvette (WYATT technology) was filled with 50 μl of DPR solution with different types of RNA, under the same conditions used in turbidity measurements. The sides of the cuvette were filled with water, and a cap was put on top. The intensity of scattered light was recorded at a scattering angle of 95° at 25°C for a period of 2 hours, collecting 7 acquisitions (8 s each). Each measurement was repeated at least 3 times. The software package DYNAMICS 7.1.9 was used for data analysis.

### Fluorescent labelling with Dylight® 488

100 µM of Dylight® 488 dye (Thermo Fisher Scientific) was used to label DPRs (PR30, GR30 and PA30) for fluorescence microscopy measurements. 50 µM of each DPR was dissolved in the reaction buffer (1x PBS, 10 mM MgCl2, 0.05% Tween 20), the dye added and then the solution incubated at room temperature for 30 min in darkness. The mixture was then desalted with a Pierce™ Peptide Desalting Spin Column (Thermo Fisher Scientific). Fluorescently labelled DPRs were protected from light and stored at -80°C.

### Fluorescent and brightfield microscopy

Microscopy measurements were carried out on a Leica DMi8 microscope. To reach appropriate fluorescence intensities, Dylight® 488-labeled DPRs were mixed with the same, but non-labelled, peptides (at a 1:200 ratio). Then, phase separation was induced by the addition of the desired amount of RNAs. The solution was incubated at 25°C and droplets were visualized with 100x magnification with brightfield, and/or fluorescence microscopy (FITC filter).

### Microscopy

The response of Neuro 2a cells to DPR and/or PSS treatment was also monitored by direct microscopic imaging. To this end, cells were plated in 24-well cell culture plates (Greiner Bio One) at a cell density of 0.1 x 106, to which DPR and/or PSS solutions at the appropriate concentrations were added the following day. After 24 hours, microscopic images were recorded on a Leica DMi8 microscope equipped with a Leica DFC7000 GT camera with a10x objective. Cell viability was evaluated by counting the cells. Data were normalized to untreated cells imaged without added compounds.

### Isothermal titration calorimetry (ITC)

ITC experiments were carried out on a MicroCal iTC200 system. For each measurement PR30, GR30, PA30 and PSS samples were dialyzed overnight into 10 mM HEPES, 50 mM NaCl, pH 7.4 buffer at various NaCl concentrations (50 mM, 150 mM or 500 mM). The PSS solution was filled in the ITC cell, the actual DPR into the syringe, from where it was injected into the cell in a series of 2 μl aliquots at 25°C, to reach the final concentrations indicated in figure legends. The raw ITC data were fitted to a single binding site model by the MicroCal PEAQ-ITC Analysis Software provided by the manufacturer.

### Microscale Thermophoresis (MST)

MST measurements were performed on a Monolith NT.115 instrument from NanoTemper accompanied with Nano Temper Control 1.1.9 software at room temperature. Premium capillaries (MonolithTM NT.115 Series) from NanoTemper company and MST-binding buffer (10 mM HEPES, pH 7.4, 150 mM NaCl, 0.05 % Tween) were used for all experiments. DPRs (PR30, GR30 and PA30) were labelled by the Protein Labeling Kit, Red-NHS, 2nd generation (Cat. No. MO-L011) - amine reactive from NanoTemper accordanceing to the protocol provided. MST data were analysed by either the NanoTemper Analysis 1.2.101 software or GraphPad Prism. In either case the data were fitted with the Hill equation, with the Hill coefficient constrained to 1, i.e., by assuming a binding stoichiometry of 1:1. Each experiment was technically repeated at least three or more times, and the mean half effective concentration (logIC50) values were calculated with standard error (SE).

### Electrophoretic mobility shift assay (EMSA)

32P-labelled mRNA from Sulfolobus acidocaldarius (5’UTR region + piece of 5’end of the ORF), with a total length of 109 ntb was prepared by 5′-end-labelling using [γ-32P]-ATP (Perkin Elmer) and T4 polynucleotide kinase (Thermo Scientific) followed by dephosphorylation using FastAP thermosensitive alkaline phosphatase from Thermo Scientific. Labelled mRNA fragments were subsequently purified using MEGAclear™ Kit (Thermo Fisher Scientific). EMSA experiments were performed with approximately 0.1 nM 32P-labelled mRNA and concentration of PR30 and GR30 ranging from 10 to 260 nM. Binding reactions of mRNA and R-DPRs and PSS_43 were prepared in buffer composed of 10 mM HEPES, 150 mM NaCl, pH 7.4 and allowed to equilibrate at 37 °C prior to electrophoresis on 6% acrylamide gels in TEB buffer (89 mM TRIS, 2.5 mM EDTA, and 89 mM boric acid).

### Molecular dynamics (MD) simulations

The all-atom molecular dynamics (MD) simulation of PSS_20 and PR10, GR10 and PA10 systems were performed with the Gromacs 2019.2 package [78–80]. The OPLS-aa force field [81] was used, as it was proved to be a right choice for both peptides [82, 83] and PSS molecule [84, 85]. Sodium and chloride ions were modelled using parameters from [86] and [87], respectively. The explicit TIP4P model was employed for water [88].

Simulations were run at three different salt concentrations (50 mM, 150 mM and 500 mM NaCl). The first step involved a single-molecule equilibration in a given salt concentration. After solvation and ion addition, the system was energy minimized and a 0.1 ns NVT equilibration was performed. The DPR peptides and the PSS molecules were then simulated in an NPT ensemble for 300 and 100 ns, respectively. For each DPR, three different initial configurations were simulated. The obtained molecule conformations were further used for the simulations of 1 PSS_20 - 1 DPR (1:1) and 7 PSS_20 - 7 DPR (7:7) systems (having 1 or 7 of both types of molecules in the simulation box). The initial configurations for these systems were constructed by placing one (for 1:1) or seven (for 7:7) copies of each final configuration of the single molecule MD simulations in a random position and orientation in a simulation box of (9 nm)3 for the 1:1 or (15.0 nm)3 for the 7:7 systems. The systems obtained were simulated in an NPT ensemble for 250 ns. The convergence of the number of contacts between PSS and peptides was used as an indicator for the system equilibration (Suppl. Fig. S12) and the analysis was done for the last 50 ns.

In all simulations, the approach described in [89] was used to control the temperature while the Parrinello–Rahman algorithm was used for the barostat [90], where the time constants were 0.5 ps and 2 ps, respectively. The temperature and the system pressure were set to 298 K and 1 bar, respectively. The long-range electrostatic interactions were calculated using the PME method [91] while the van der Waals interactions were described using the Lennard-Jones potential and a 1.0 nm cut-off. LINCS [92] algorithm constrain the bonds between H and heavy atoms in the PSS and peptide molecules, while for water molecules the SETTLE algorithm [93] was used. A 2-fs time step was used for integrating the equations of motion. VMD software has been used for visualizations [94].

### NMR

PSS-DPR interactions were also monitored by nuclear magnetic resonance (NMR) on a Varian INOVA 800 MHz spectrometer equipped with a Bruker console and cryo-probe head. Data processing was performed using Topspin v. 4, and data was plotted by GraphPad Prism and Topspin v.4. The sample buffer was 50 mM sodium phosphate, 50 mM NaCl, pH 7.0 and 5% D_2_O.

### Cell culture

Neuro-2a cel were obtained from cell line service (CLS), and U2OS cell lines were maintained in Dulbecco’s modified Eagle’s medium (DMEM, Gibco) supplemented with 10% FBS and 1% penicillin/streptavidin (Gibco) at 37°C in a humidified incubator with 10% CO_2_.

### Assaying cellular ATP content

To assess changes in cell viability, we measured the ATP content of Neuro 2a cells. To this end, cells were seeded in white transparent-bottom 96 well microplates from life sciences research (Perkin Elmer) at 20,000 cells per well and incubated at 37°C with 5% CO_2_ overnight. The medium was carefully removed, PSS at the appropriate concentration was added to the wells, and the plate was then incubated for 24 hours. The ATP content indicative of cell viability was measured at various time points by ATPlite Luminescence Assay System (PerkinELmer). Luminescence was measured by the BioTek SynergyTM Mx plate reader. The luminescence of cells without added compounds was taken as 100%.

### iPSC-derived motor neurons

iPSCs were purchased from Sigma-Aldrich (iPSC Epithelial-1, IPSC0028). iPSC colonies were kept in Essential 8 flex medium (Gibco), supplemented with penicillin-streptomycin (1000 U/ml) on Geltrex™ Reduced-Growth Factor Basement-Membrane Matrix, LDEV-free (Gibco) coated plates incubated at 37°C and 5% CO_2_. Upon reaching 80 to 90% confluency, they were passaged using 0.5 mM Promega EDTA (Thermo Fisher Scientific) diluted in Dulbecco’s Phosphate-Buffered Saline (Gibco). The differentiation of iPSCs into motor neurons was performed as earlier described [95].

### Mitochondrial transport measurements in iPSC-derived motor neurons

To determine the effect of R-DPRs and PSS on axonal transport phenotypes in motor neurons, we performed mitochondrial transport measurements as previously described [64]. Briefly, motor neurons were seeded in PenoPlate 96-well (Revvity) at a density of 10,000 cells per well and matured to day 40. Motor neurons were pre-treated with 0.2 µM GR30 for 8h (or dH_2_O for control), after which the media of the cells was replaced by media containing 0µM, 5µM or 10µM of PSS 22 and left on overnight. Next, the cells were washed with BrainPhys^TM^ Without Phenol Red (Fisher Scientific) and stained with MitoTracker^TM^ Green diluted in BrainPhys^TM^. Measurements were performed on a ZEISS Celldiscoverer 7 microscope with a x40 air lens (live imaging system at 37°C and with 5% CO_2_) at LiMoNe (VIB-KU Leuven). For each data point, 200 sequential images were acquired at a rate of one frame per second (1 Hz), and image analysis was performed in Fiji using the automated tracking plugin TrackMate. To avoid unconscious bias, the analysis was carried out by an experimenter blinded for the treatment. Data were analyzed and visualized using GraphPad Prism 10.1.1. For statistical analysis, three independent biological replicates were used.

### Mouse toxicity

Mouse toxicity analysis was carried out by Neuro-Sys Vivo SAS (Gardanne, France). The aim of the study was to administer PSS_22 in the brain of 12-week-old adult male C57BL/6 mice (by intracerebroventricular (ICV) injection), to evaluate its potential acute toxicity. Mice were monitored (body mass, daily observation) for 7 days.

PSS_22 was injected in 5 μl volume in sterile PBS as carrier, at either 1 mM or 10 mM concentration (5 mice each), and a toxicity score was given based on assessing: seizure, atypical behavior, hyperactivity, normal grooming/eating.

## Abbreviations

ALS: amyotrophic lateral sclerosis
APS: ammonium persulfate
ASO: antisense oligonucleotide
BBB: blood-brain barrier
BLI: bio-layer interferometry
DAPI: 4,6-diamidino2-phenylindole
DLS: dynamic light scattering
DPR: dipeptide repeat
EMSA: electrophoretic mobility shift assay
fALS: familial amyotrophic lateral sclerosis
FDA: Food and Drug Administration
FRAP: fluorescence recovery after photobleaching
FTD: frontotemporal dementia
FUS: fused in sarcoma
G3BP1/2: Ras GTPase-activating protein-binding protein ½
HRE: hexanucleotide (G4C2) repeat expansion
IDP: intrinsically disordered protein
ITC: isothermal titration calorimetry
LLPS: liquid-liquid phase separation
MD: molecular dynamics
MST: microscale thermophoresis
MNs: motor neurons
NHS: N-Hydroxysuccinimide
NPM1: Nucleophosmin
PBS: phosphate-buffered saline
PEG: polyethylene glycol
PSS: polystyrene sulfonate
PSS_Rhod: Rhodamine-labelled heterogenous mixture of PSS
RAN: repeat-associated non-AUG translation
Rg: radius of gyration
R-DPR: arginine-rich DPR
RDF: radial distribution function
RRM: RNA-recognition motif
sALS: sporadic amyotrophic lateral sclerosis
SCA36: spinocerebellar ataxia type 36
SG: stress granule
SV2: vesicle 2 protein
TDP-43: TAR DNA-binding protein 43
TEMED: N,N,N′,N′-tetramethyl ethylenediamine
ThS: thioflavin S
ThT: thioflavin T
XDP: X- linked dystonia Parkinsonism

## Acknowledgements

This work was supported by grants K124670, K131702 and RGH_24 (RGH 151464) grant agreement with the National Research, Development and Innovation Office, Hungary (NKFIH). Funds are also acknowledged to the National Science Centre, Poland (grant no. 2018/31/D/ST5/01866) (P.B.) and grant no 2021/43/B/ST8/01900 by the National Science Center, Poland, Opus Project, for A.B-S. The research of L.V.D.B. was supported by VIB, KU Leuven (C14/22/132, IDN/22/ 012 and “Opening the Future” Fund), the “Fund for Scientific Research Flanders” (FWO-Vlaanderen; G0C1620N, G088523N and G026924N), Target ALS, the ALS Liga België (A Cure for ALS), the Generet Award for Rare Diseases and the Chair Dr. Helen Camerlynck. The contribution of an FWO PhD fellowship in strategic basic research (FWOSB77, to JA) is also acknowledged. TL was holder of a postdoctoral innovation mandate (grant no. HBC.2022.0194) by the Flanders Innovation & Entrepreneurship Agency (VLAIO). We also thank the computational resources and services used in this work that were provided by the VSC (Flemish Supercomputer Center), funded by the Research Foundation - Flanders (FWO) and the Flemish Government, and Poland’s high-performance computing infrastructure PLGrid (HPC Centers: ACK Cyfronet AGH) within computational grant no. PLG/2023/016229. The authors are also indebted to Rani Baes (Microbiology, DBIT, VUB) for the RNA constructs provided for the EMSA experiments.

## Supplementary material

**Fig. S1.**
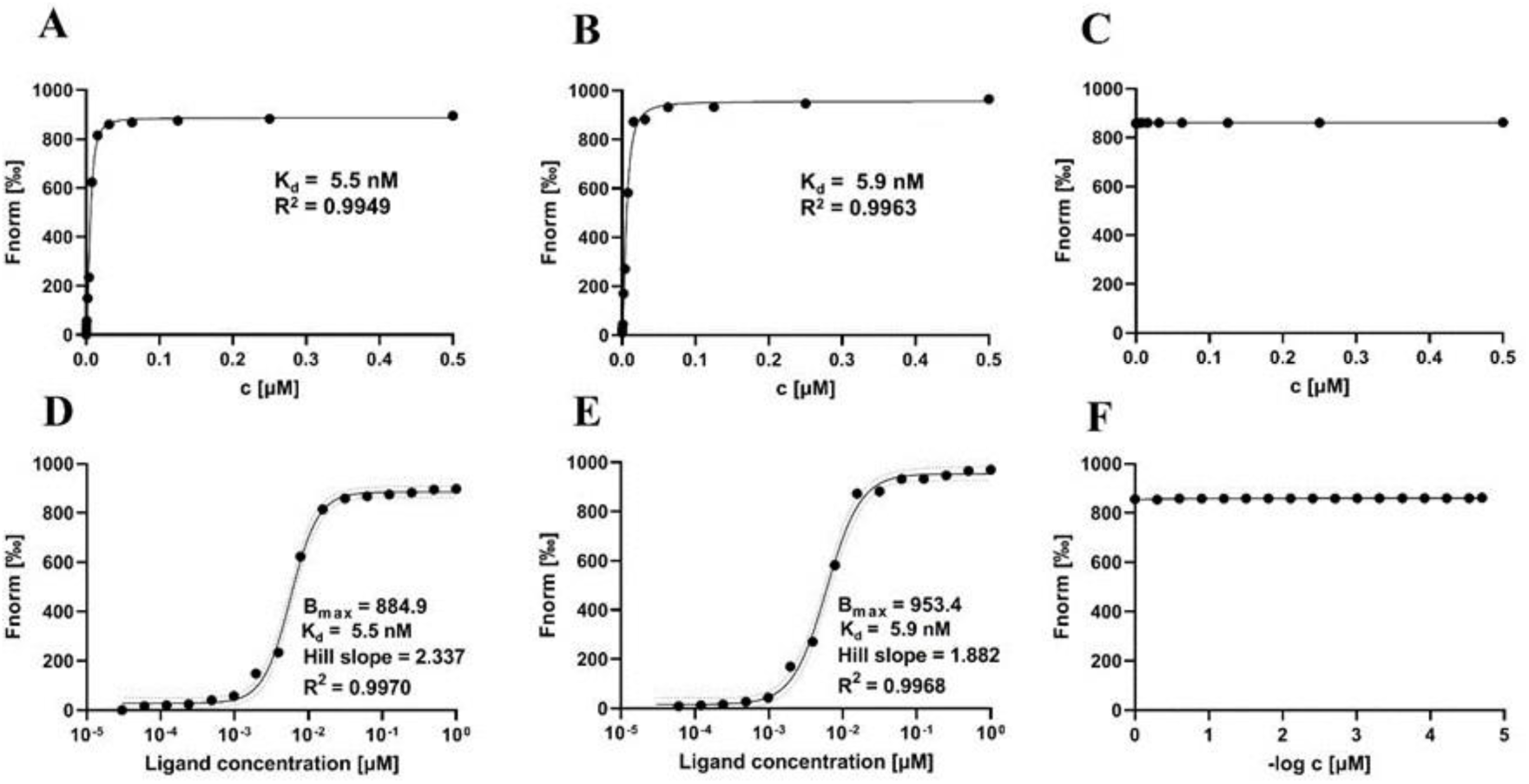
Tight binding between DPRs and PSS_43. Binding was measured by MST (L-R), the binding curves of PSS_43 and PR30 (L,O), GR30 (M,P), and PR30 (N,R) in a buffer of 10 mM HEPES, 150 mM NaCl, 0.05 % Tween are shown. The concentration of DPRs (labelled with Red-N-Hydroxysuccinimide (Red-NHS), 2nd generation dye from Nanotemper) was 2 µM, while concentration of PSS_43 varied from 0.03 nM to 500 nM. Measurements were repeated three times and data points are averaged with SEM, maximum fluorescence (Bmax) and R2. For all MST curves, apparent IC_50_ values were calculated using the Hill-equation.

**Figure S2.**
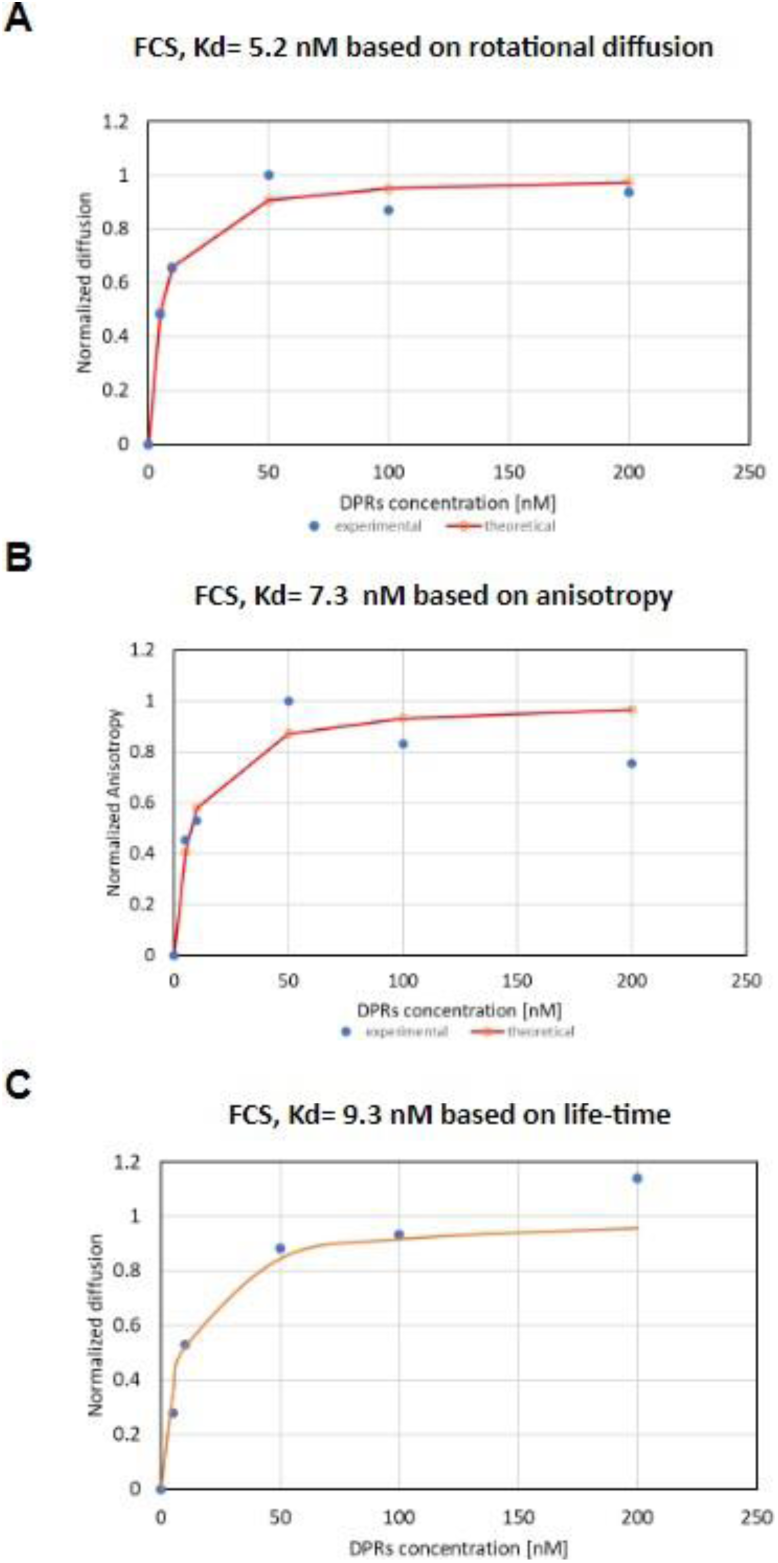
Single-molecule fluorescence measurement of PSS_43 - PR30 binding. Kd of PSS_43 – PR30 binding was determined by single-molecule fluorescence, based on measuring rotational diffusion (A), anisotropy (B) and fluorescence lifetime (C) of DyleLight 488-labeled PR30.

**Figure S3.**
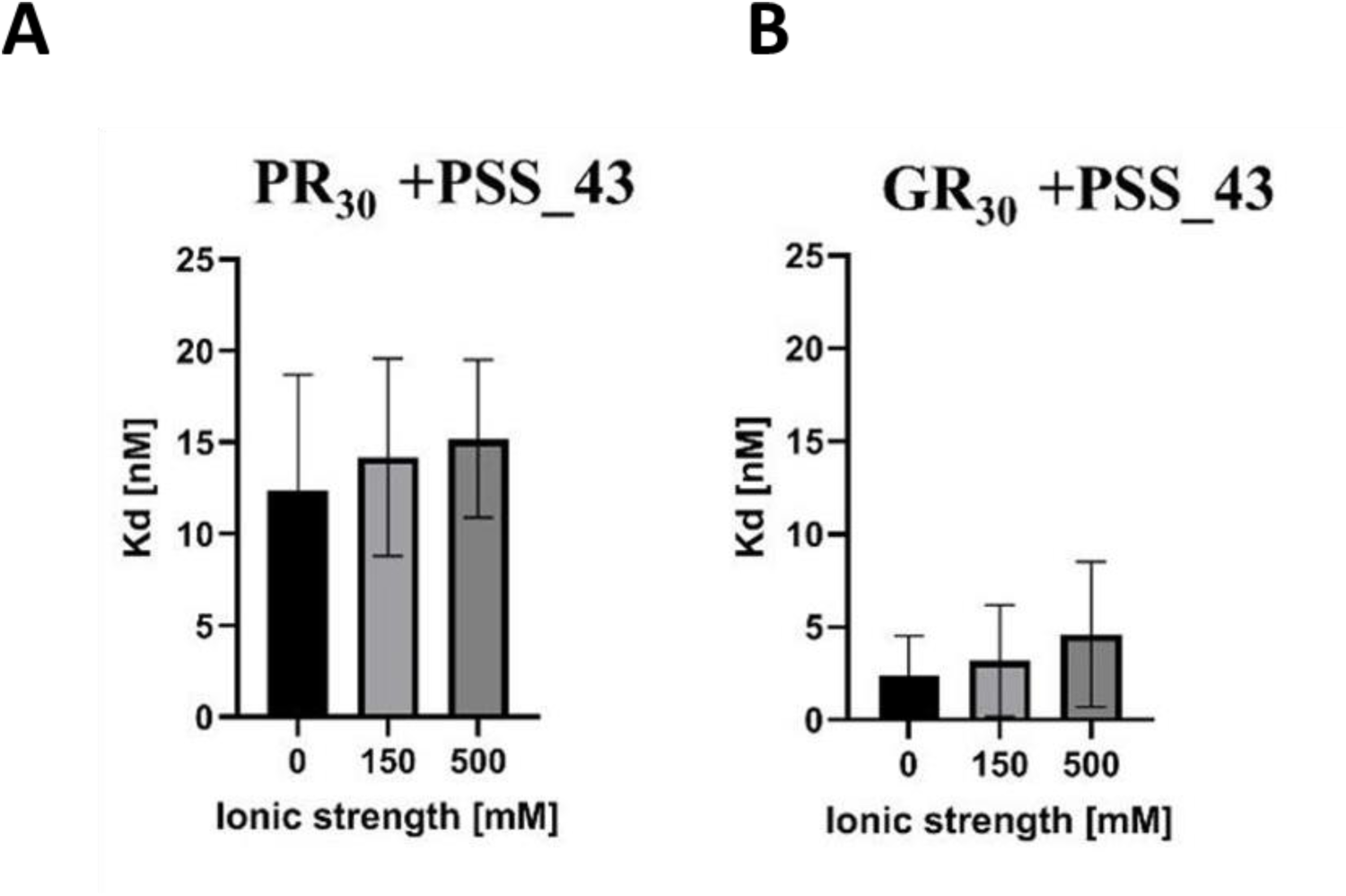
Salt titration of PSS - R-DPR systems. PR30 and GR30 were titrated by PSS_43, and Kd of interaction measured by ITC, at various salt concentrations (0 mM, 150 mM and 500 mM).

**Figure S4.**
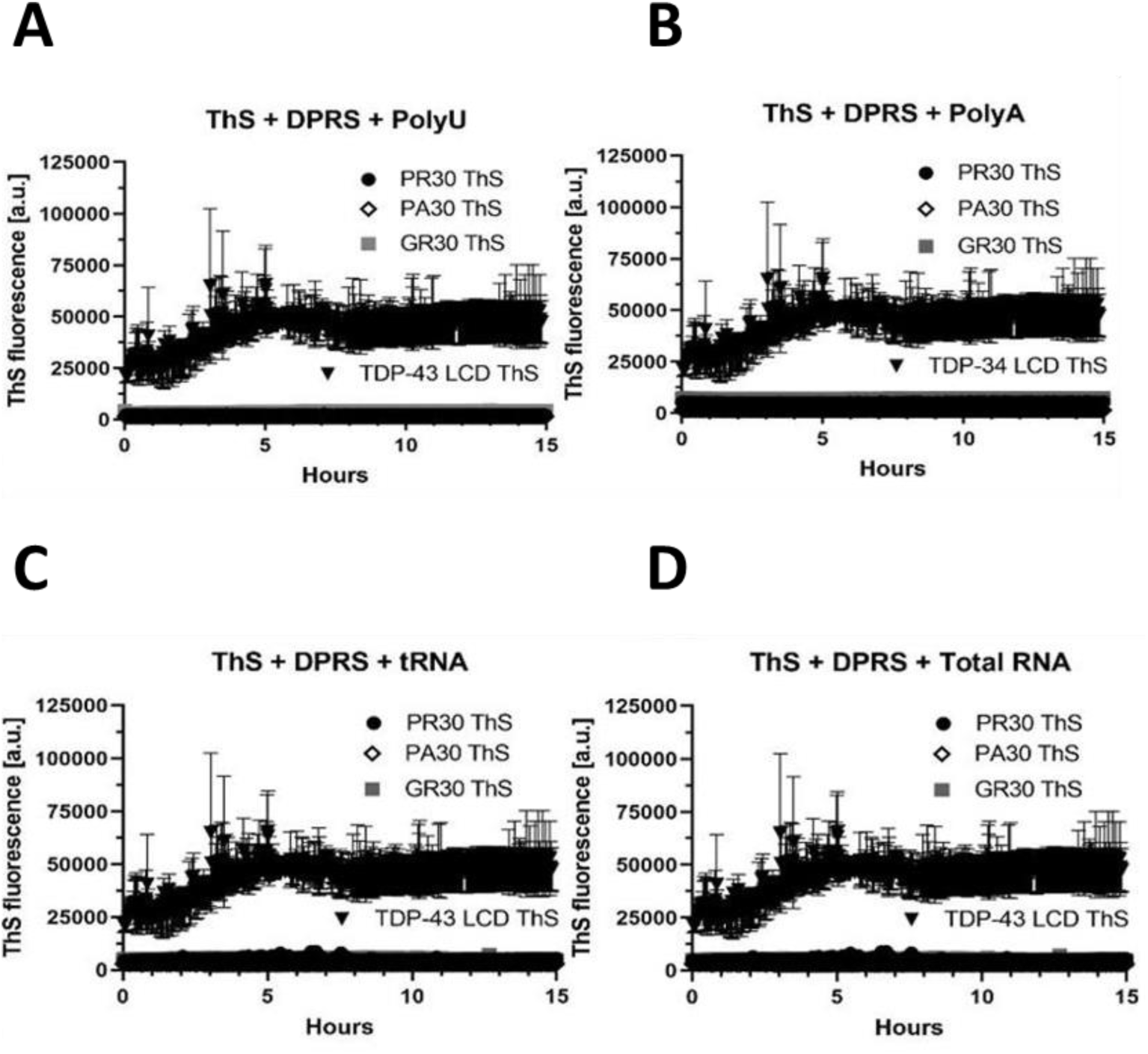
ThT and ThS curves of R-DPR - RNA systems. LLSP of R-DPRs was initiated by the RNA indicated and followed over a time of 15 h, in the presence of ThS, to indicate the possible transition to an aggregated state

**Figure S5.**
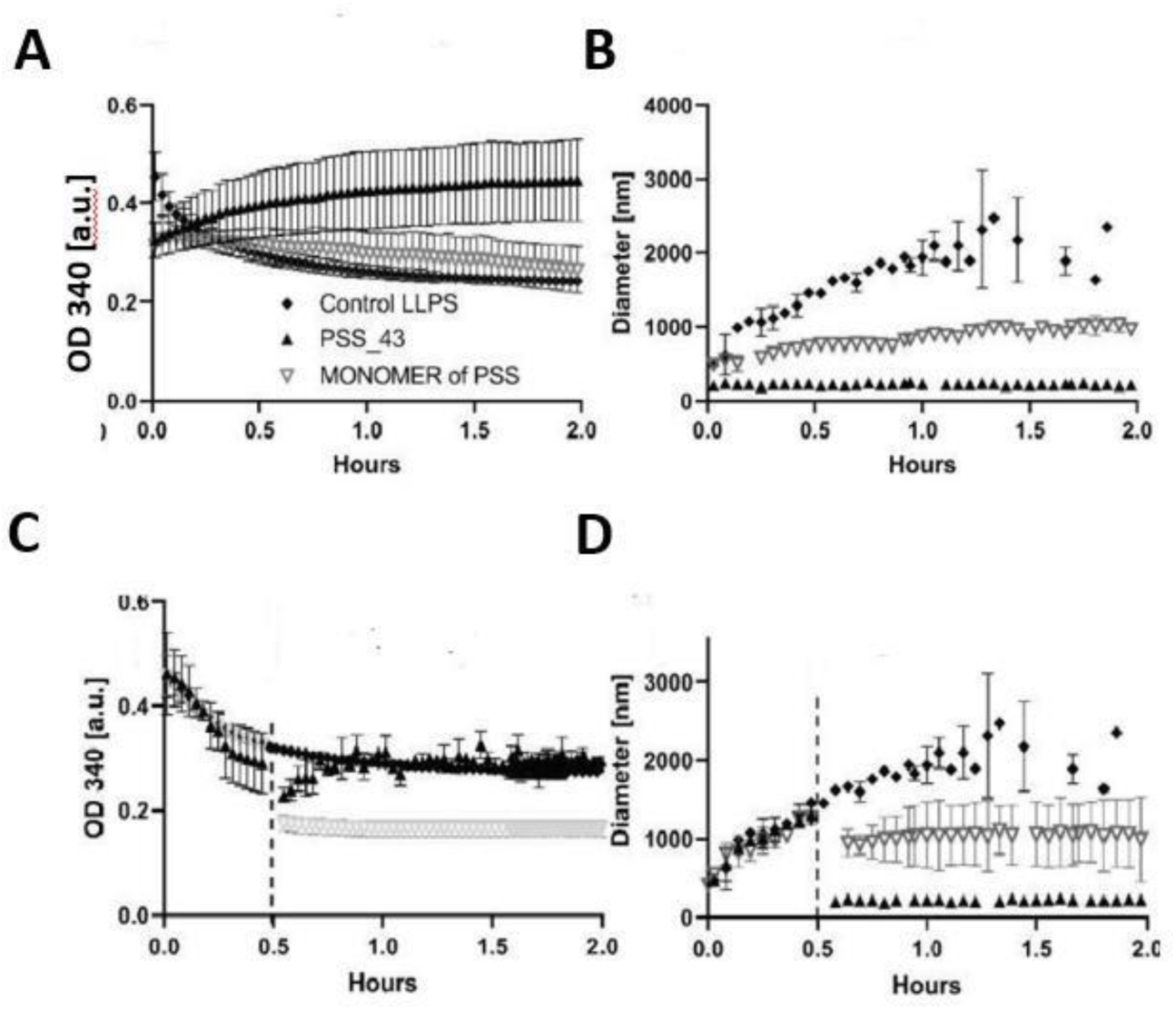
Effect of PSS_43 on GR30 LLPS. LLPS of GR30 was initiated by polyU in the presence of PSS_43 or the monomer of PSS, and monitored by adding PSS before (A, B) or 30 min after (C, D) LLPS. The LLPS was followed by turbidity (OD340, A, C) or DLS (B, D). The experiments were performed at 150 mM NaCl, pH 7.4, at 50 µM PR30 and 0.5 µg/µl RNA concentration. The molar ratio of PSS_43 and the monomeric version was 1:43, which corresponds to the number of styrene sulfonate units in PSS_43.

**Figure S6.**
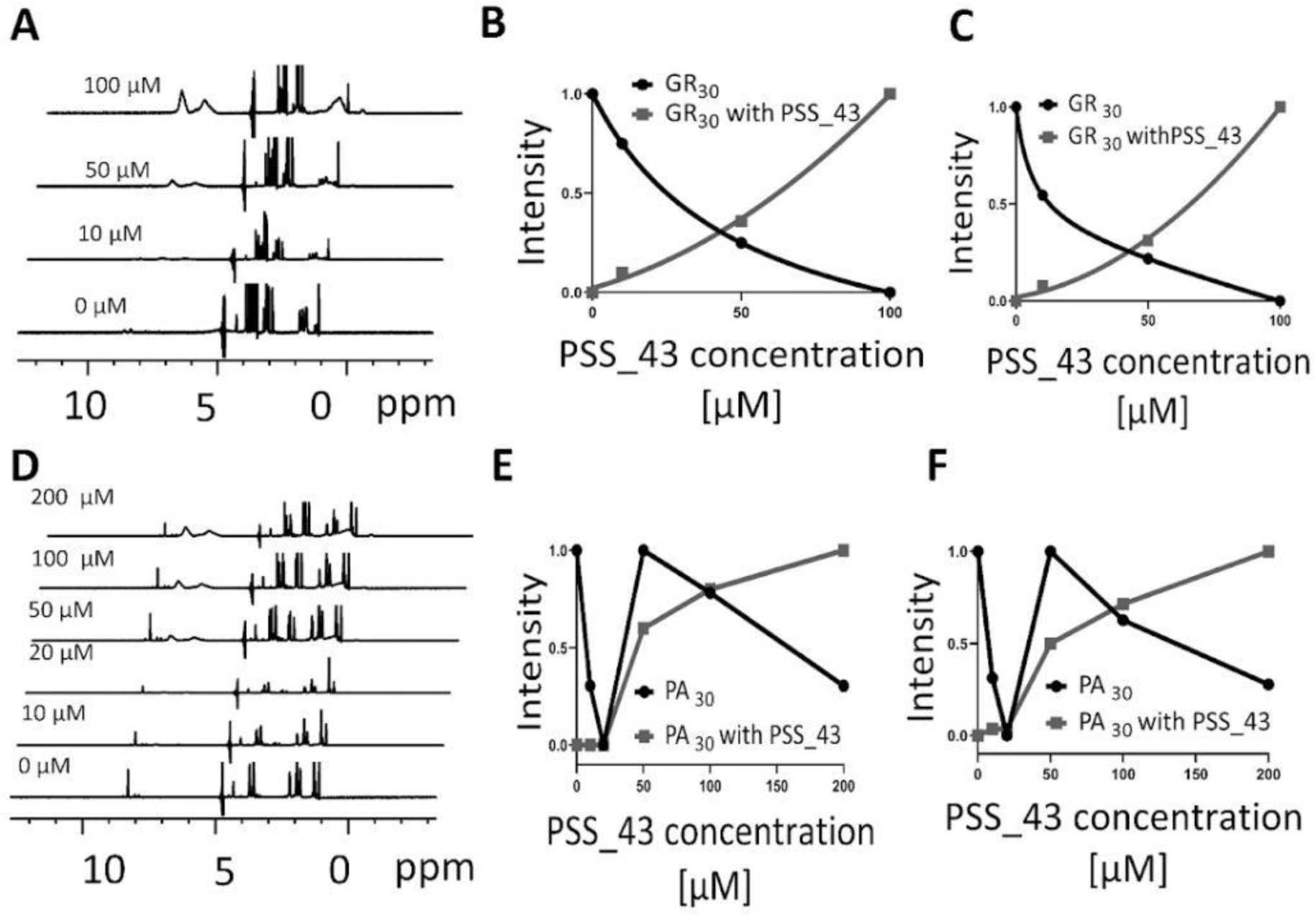
NMR of PSS - GR30 and PSS - PA30 systems. ^1^D^1^H NMR spectrum of DPR PR30 (A) and PA30 (D) was recorded in the presence of increasing PSS_43 concentrations (up to 300 μM). Peak intensities in the aromatic and amide (6-9 ppm) and aliphatic (1-3 ppm) regions were integrated and plotted separately for the DPR and PSS as a function of PSS concentration for GR30 (B, C) and PA30 (E, F).

**Figure S7.**
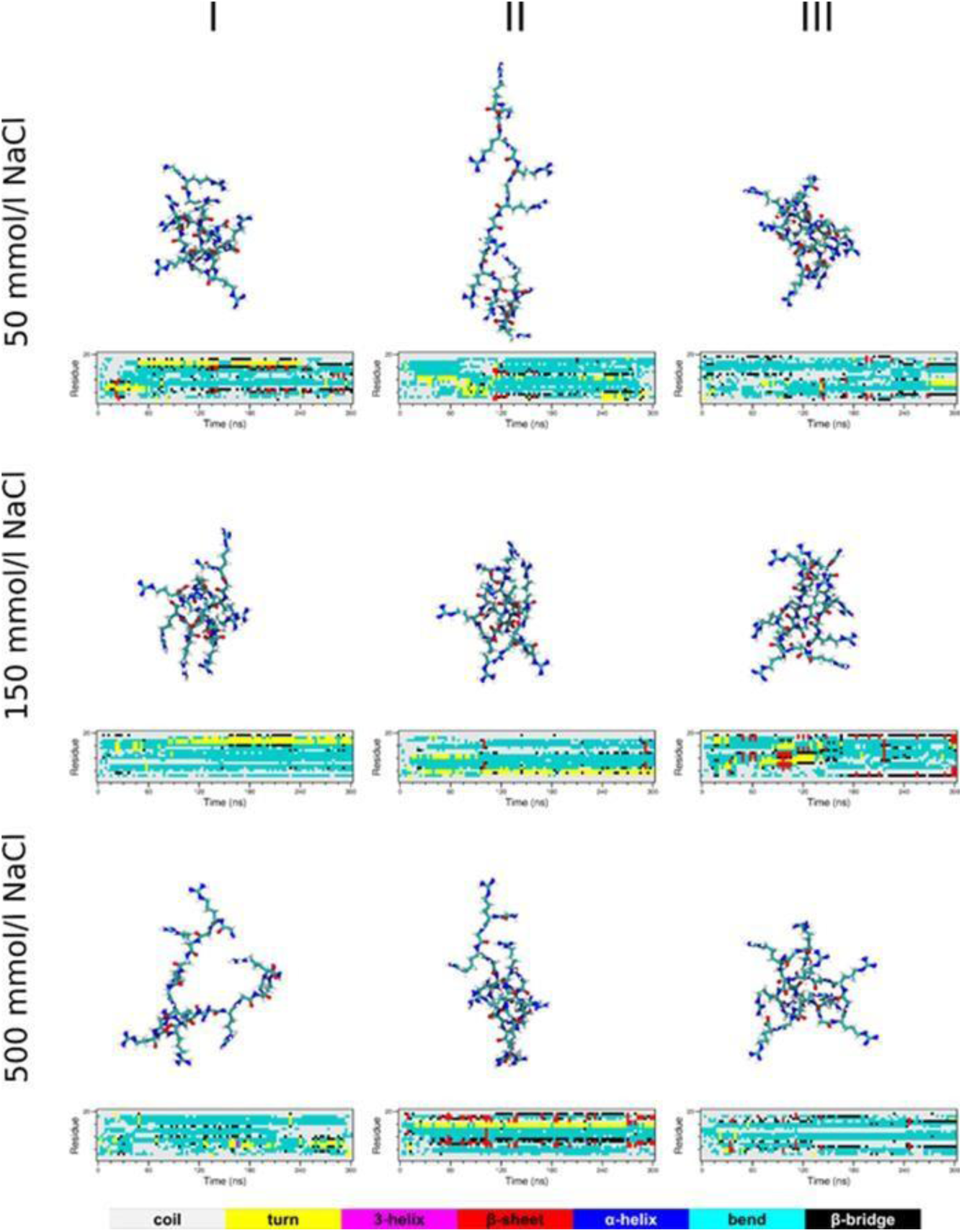
GR10 peptide conformations in MD simulations. Representative conformations are shown after 300 ns MD simulations and its secondary structure evolution at various NaCl concentrations. Results for the three different initial configurations (I, II, and III) are presented in columns.

**Figure S8.**
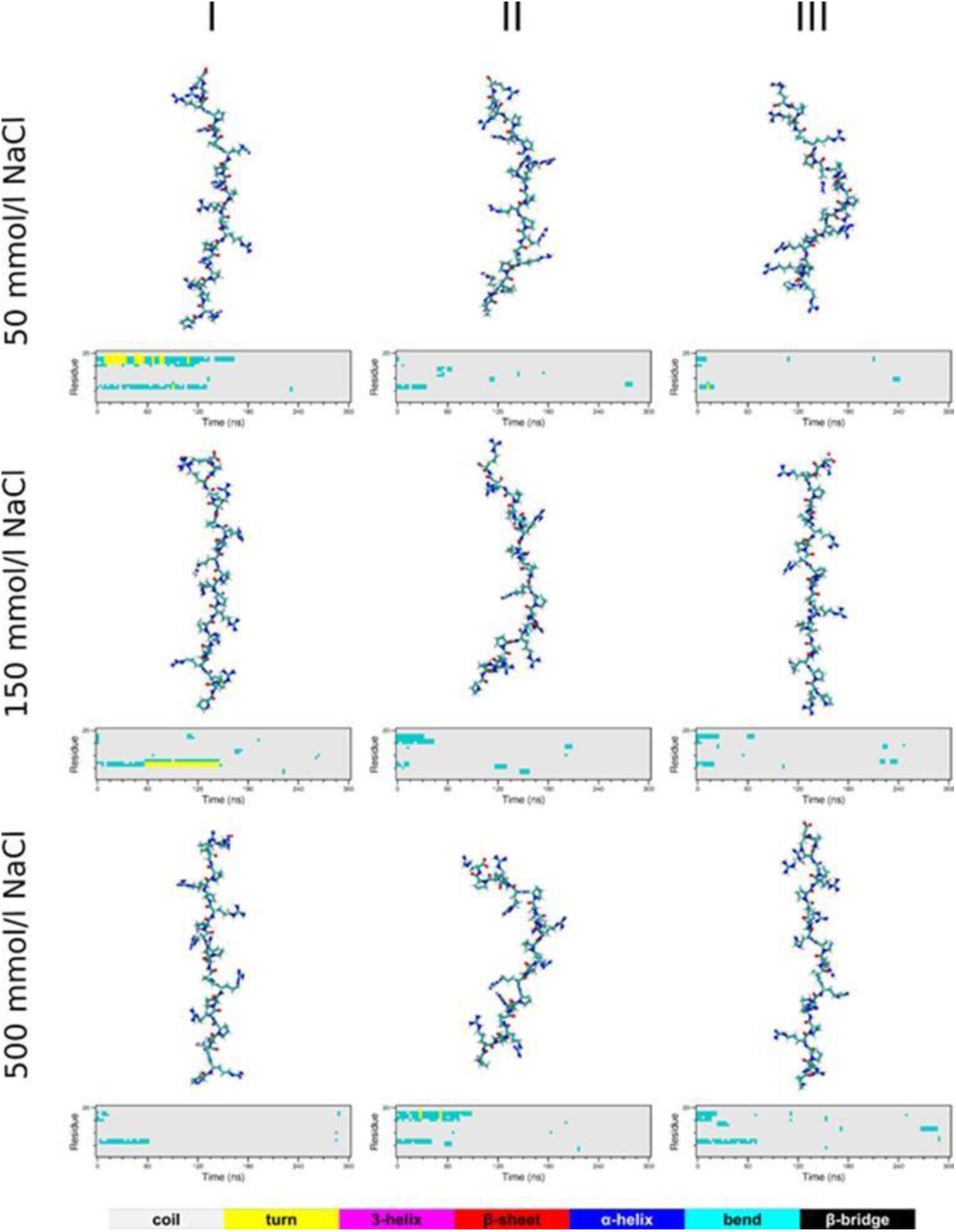
PR10 peptide conformations in MD simulation. Representative conformations after 300 ns MD simulations and its secondary structure evolution at various NaCl concentrations. Results for the three different initial configurations (I, II, and III) are presented in columns.

**Figure S9.**
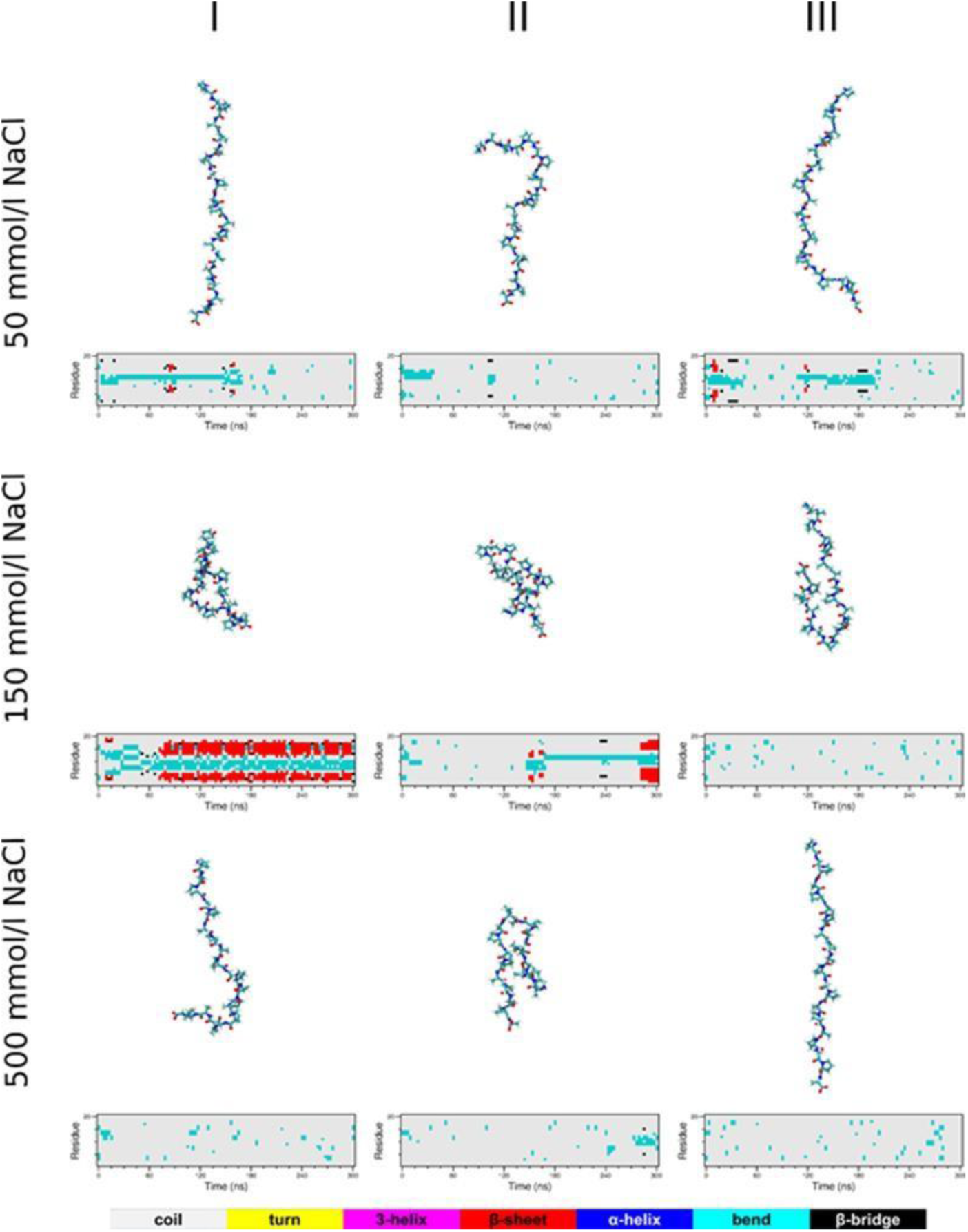
PA10 peptide conformations in MD simulations. Representative conformations after 300 ns MD simulations and its secondary structure evolution at various NaCl concentrations. Results for the three different initial configurations (I, II, and III) are presented in columns.

**Figure S10.**
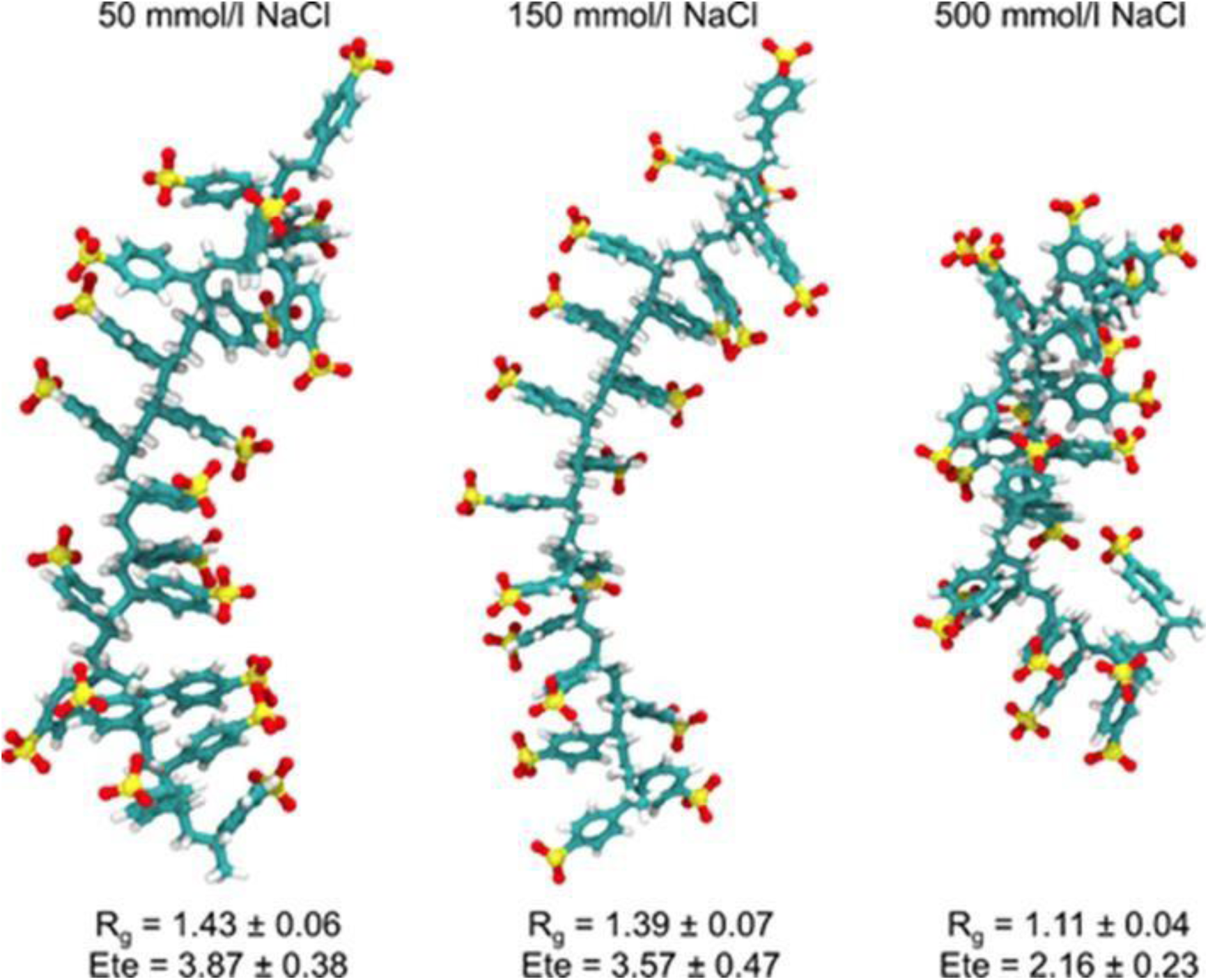
PSS20 conformations in MD simulation. Representative conformations after 100 ns MD simulations at various NaCl concentrations, along with their radius of gyrations (Rg) and end-to-end distances (Ete), averaged over the last 50 ns.

**Figure S11.**
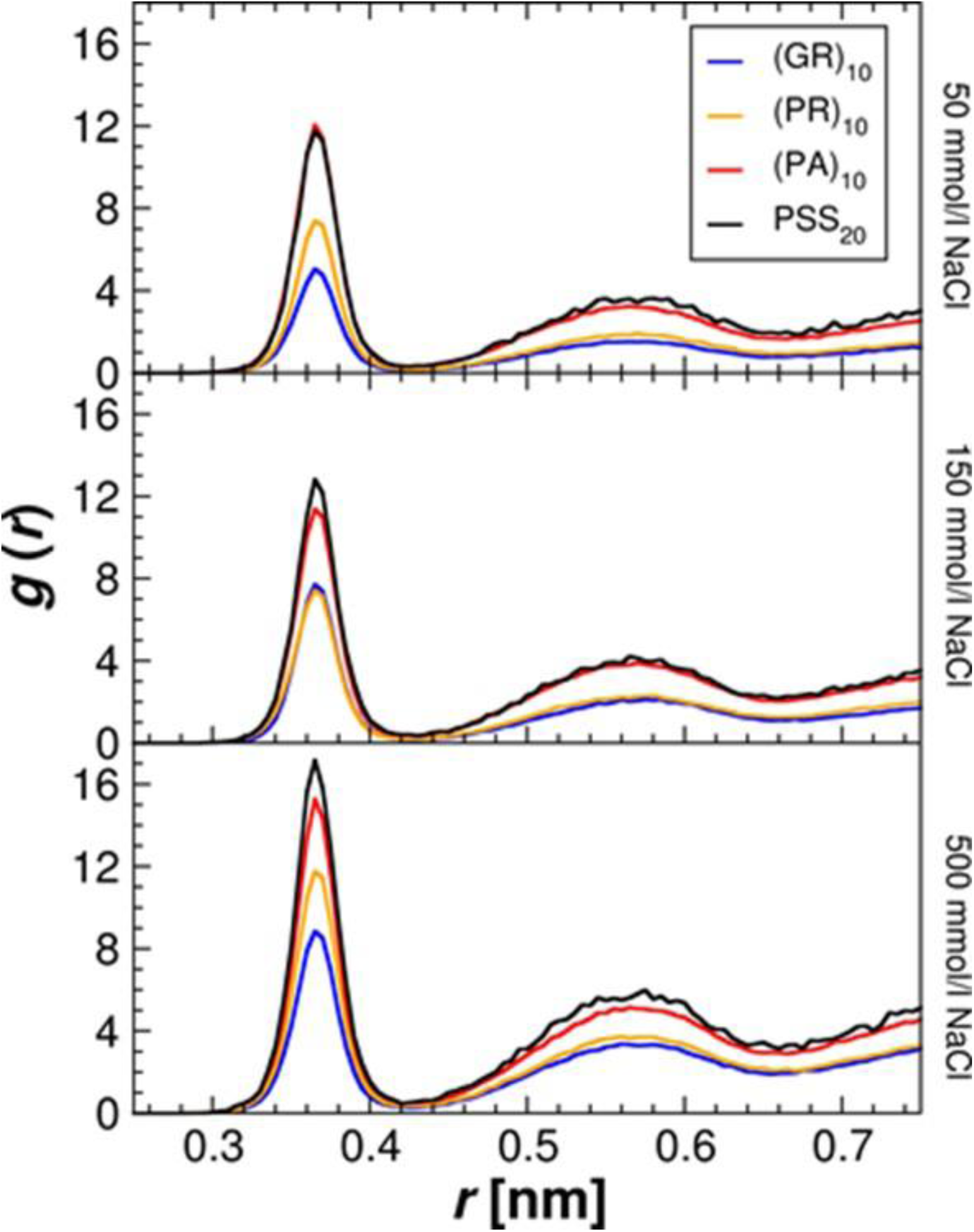
The radial distribution function (RDF) of trajectories of MD simulations. MD simulations of GR10, PR10, P!10 and PSS_20 were analyzed for RDF, as defined between the S atom from PSS and Na+ counterion in the investigated 1:1 PSS20-Peptide systems at various NaCl concentrations.

**Figure S12.**
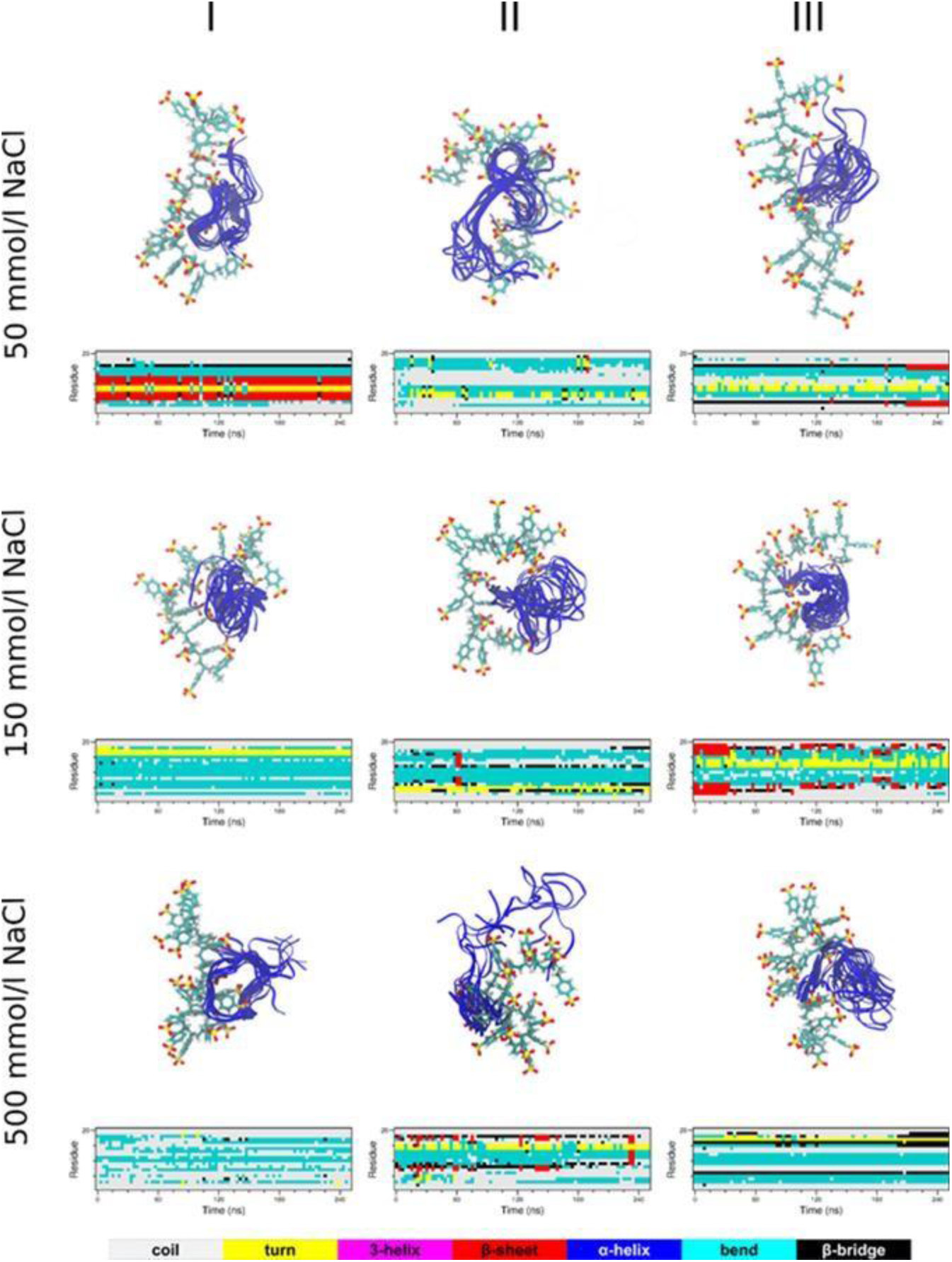
MD simulation snapshots of GR10-PSS20 systems. Representative conformations are shown of trajectories at various NaCl concentrations, aligned to the PSS20 molecule. GR10 conformations, saved every 20 ns, are shown as blue ribbons. The secondary structure evolution of GR10 is shown below the snapshots. Results for the three different initial configurations (I, II, and III) are presented in columns.

**Figure S13.**
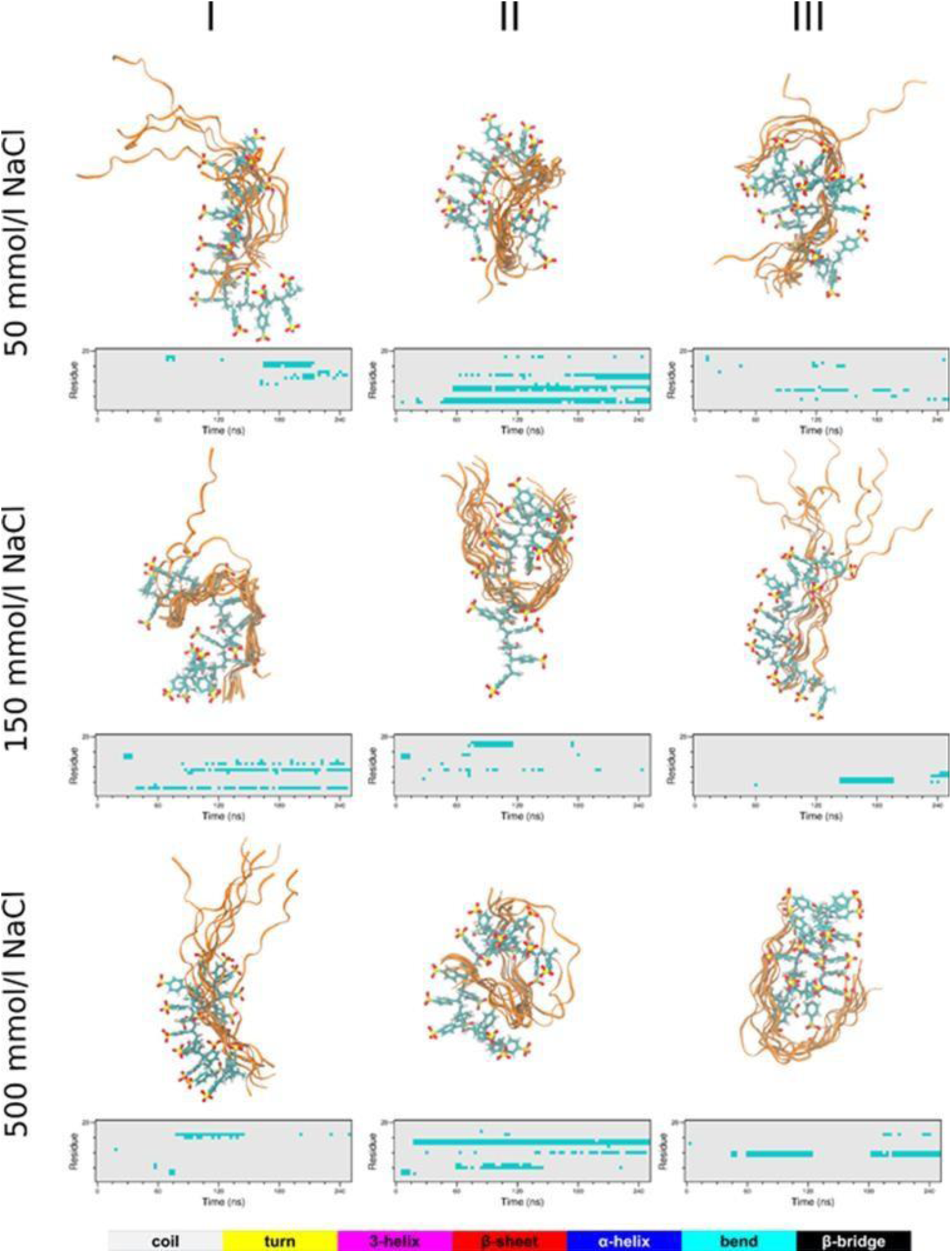
MD simulation snapshots of PR10-PSS20 systems. Representative conformations are shown of trajectories at various NaCl concentrations, aligned to the PSS20 molecule. PR10 conformations, saved every 20 ns, are shown as orange ribbons. The secondary structure evolution of PR10 is shown below the snapshots. Results for the three different initial configurations (I, II, and III) are presented in columns.

**Figure S14.**
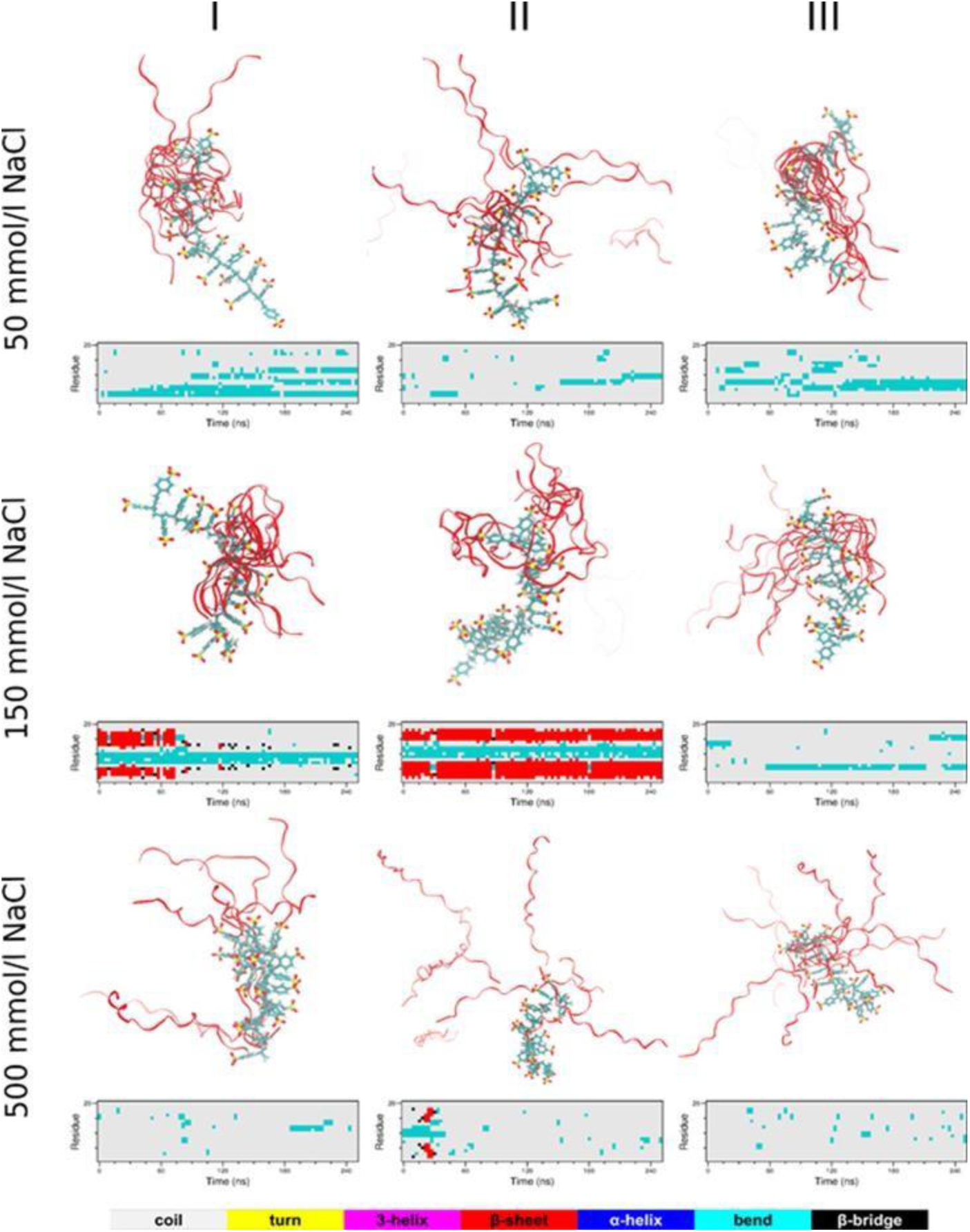
MD simulation snapshots of PA10-PSS20 systems. Representative conformations are shown of trajectories at various NaCl concentrations, aligned to the PSS20 molecule. PA10 conformations, saved every 20 ns, are shown as red ribbons. The secondary structure evolution of PA10 is shown below the snapshots. Results for the three different initial configurations (I, II, and III) are presented in columns.

**Figure S15.**
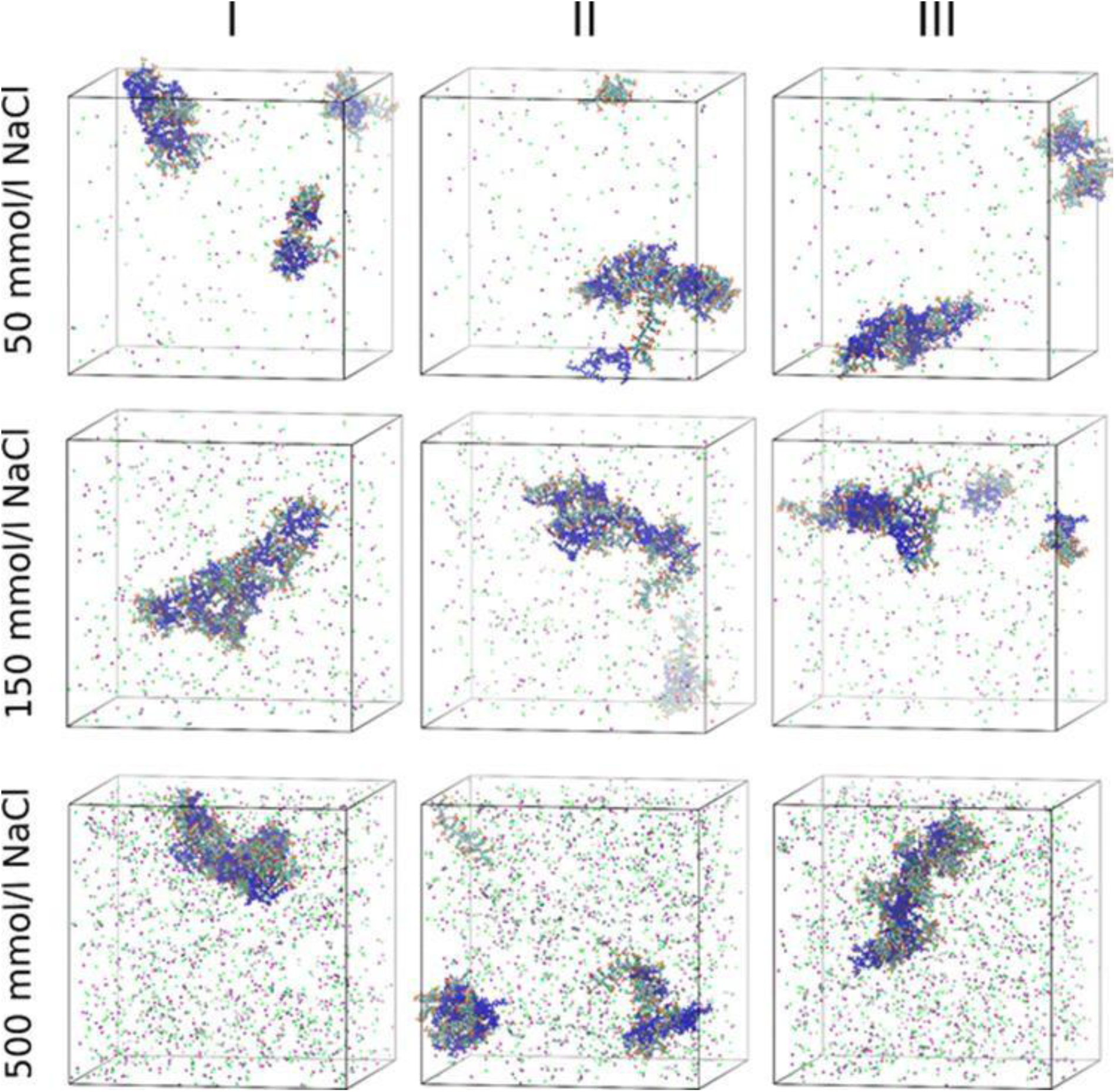
MD simulation snapshots of 7PSS20-7GR10 systems. Representative conformations are shown of trajectories at various NaCl concentrations. GR10 molecules are highlighted in blue. The Na+ and Cl- ions are highlighted as green and pink spheres, respectively. Results for the three different initial configurations (I, II, and III) are presented in columns.

**Figure S16.**
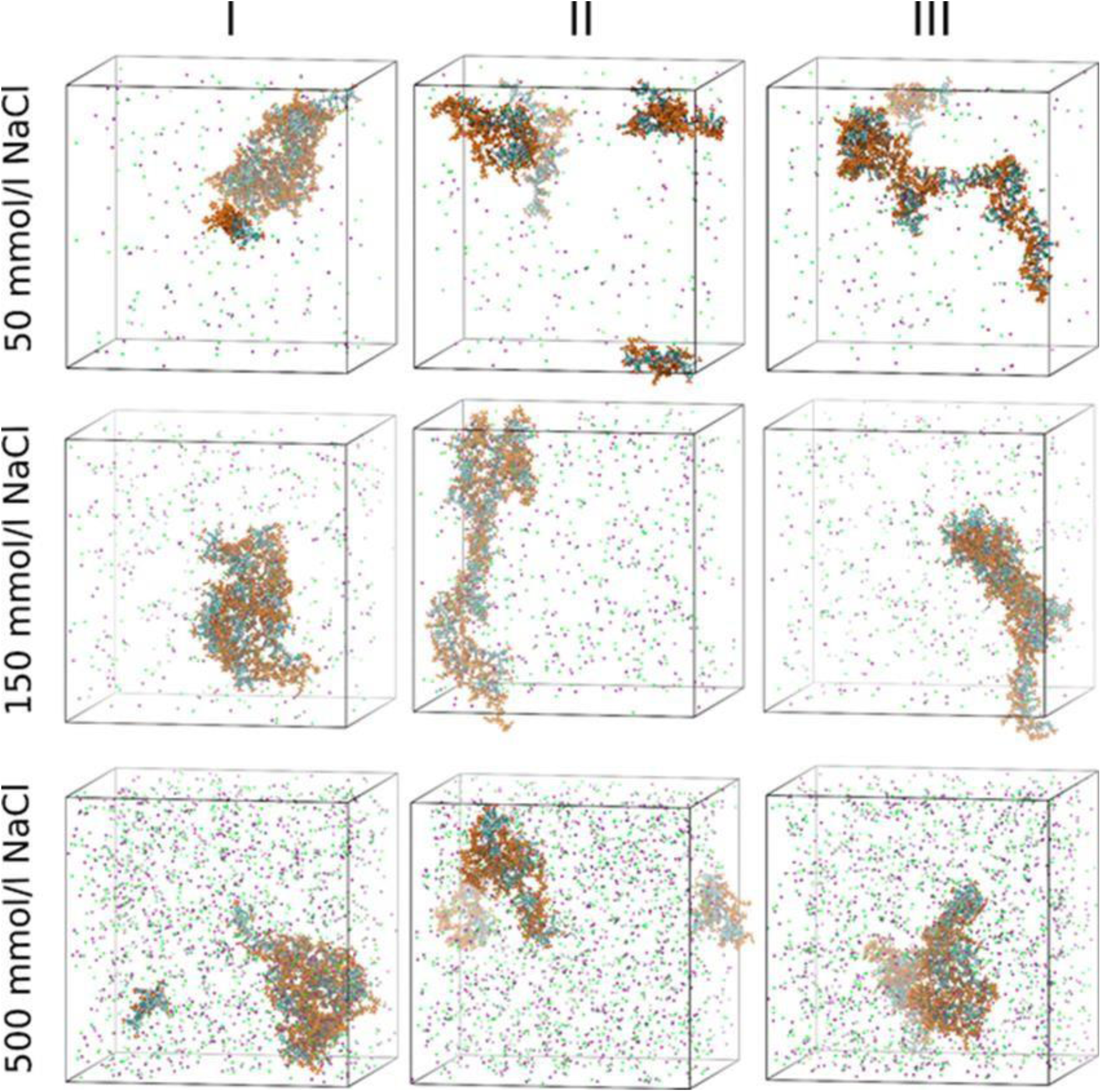
MD simulation snapshots of 7PSS20-7PR10 systems. Representative conformations are shown of trajectories at various NaCl concentrations. PR10 molecules are highlighted in orange. The Na+ and Cl- ions are highlighted as green and pink spheres, respectively. Results for the three different initial configurations (I, II, and III) are presented in columns.

**Figure S17.**
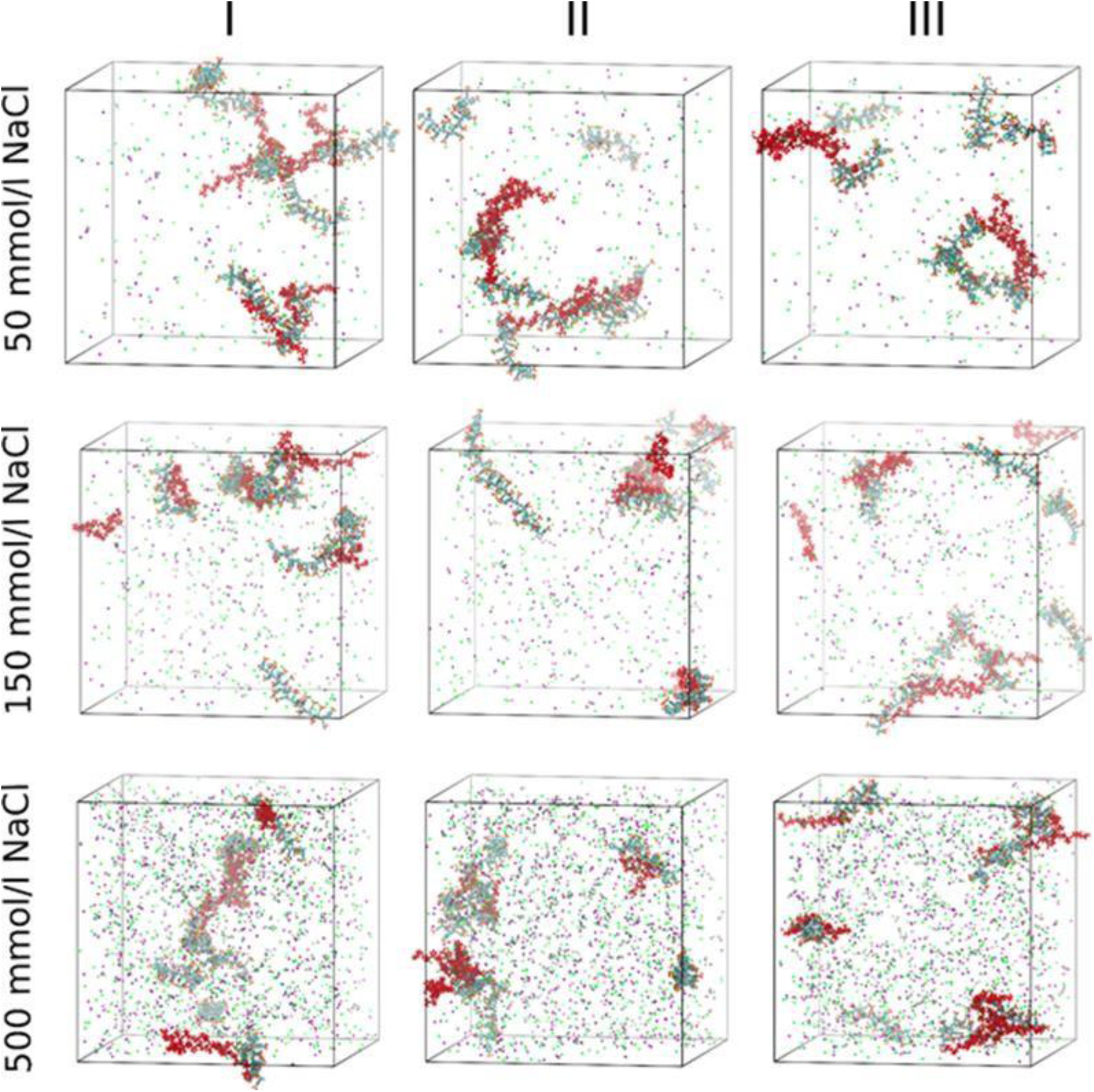
MD simulation snapshots of 7PSS20-7PA10 systems. Representative conformations are shown of trajectories at various NaCl concentrations. PA10 molecules are highlighted in red. The Na+ and Cl- ions are highlighted as green and pink spheres, respectively. Results for the three different initial configurations (I, II, and III) are presented in columns.

**Figure S18.**
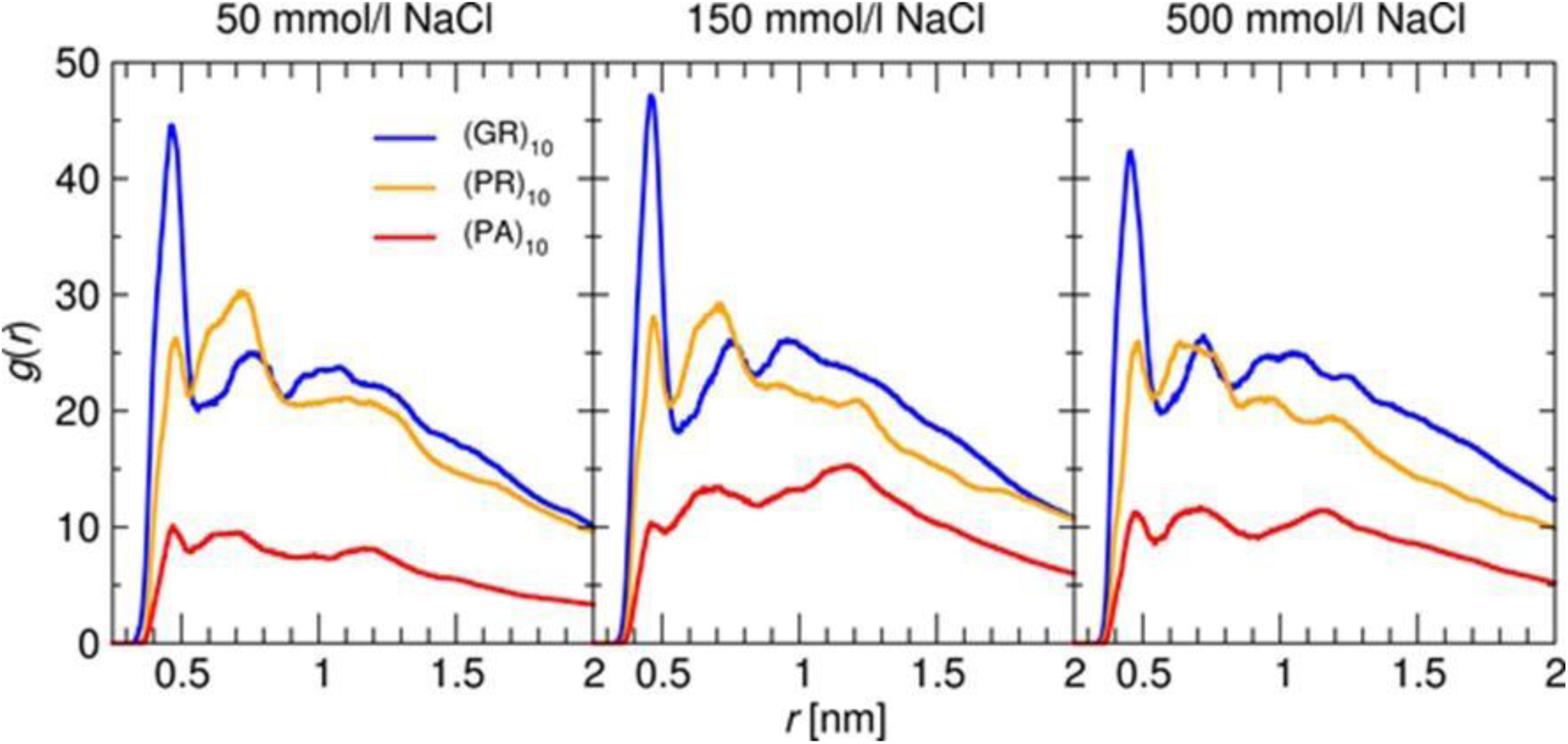
The radial distribution function (RDF) between the Cα atoms and S atom. MD simulations of 7 PSS_20 : 7 Peptide systems were run at various NaCl concentrations, averaged over three different initial configurations and analyzed for RDF.

**Figure S19.**
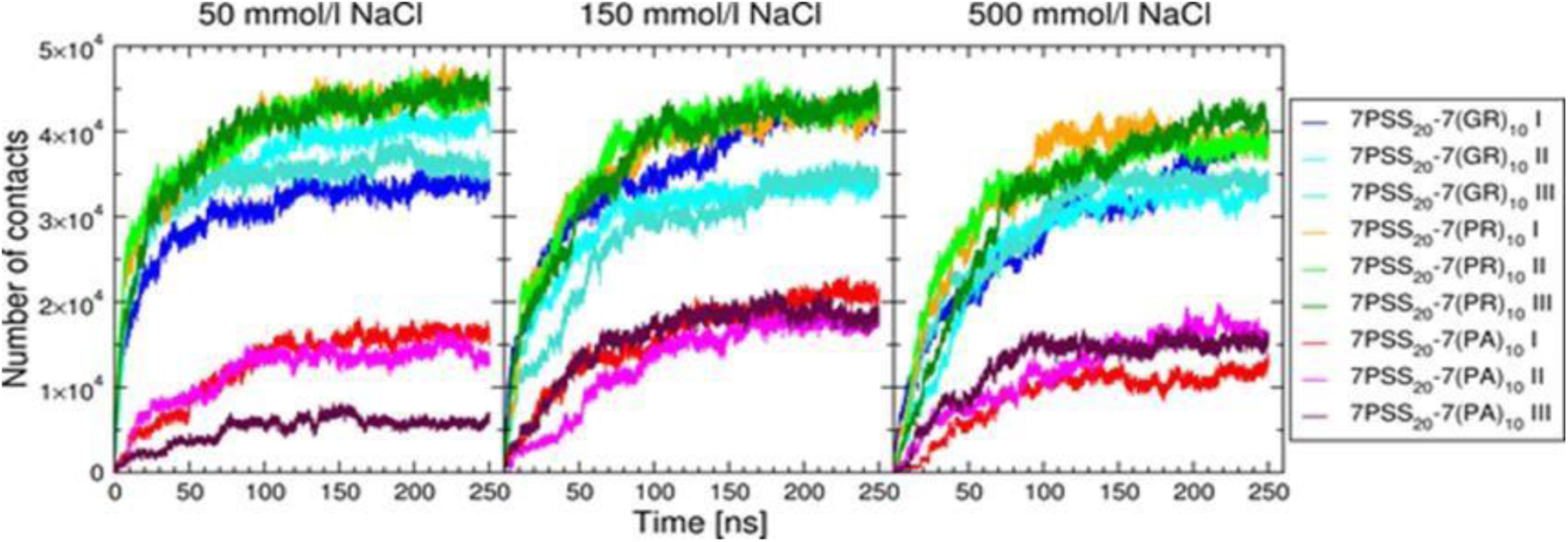
Evolution of the number of contacts. Evolution of the number of contacts between PSS_20 and R-DPR molecules in 7:7 systems simulated, as a function of the simulation time for systems at various NaCl concentrations.

**Table S1.**
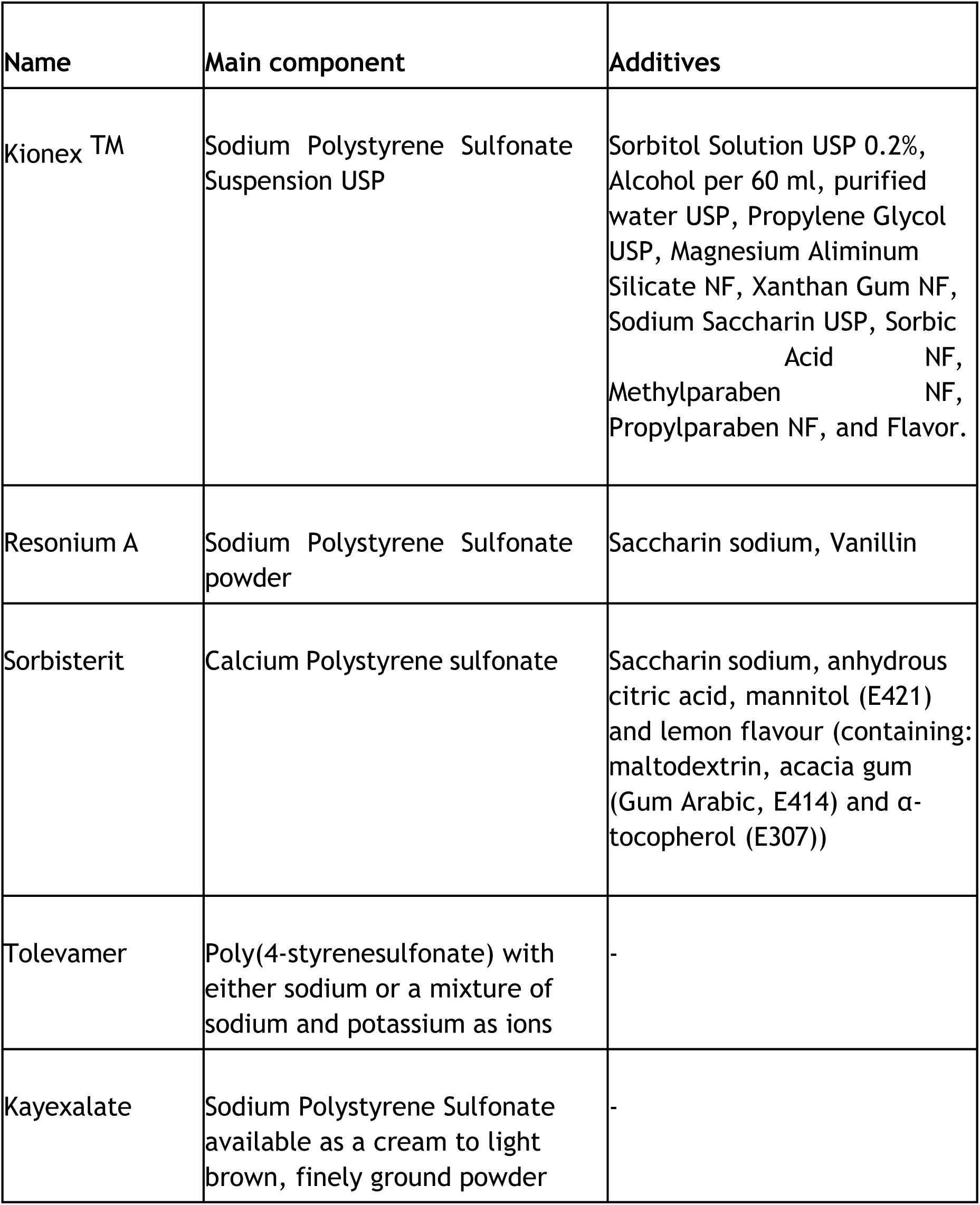
Drugs on the market that contain polystyrene sulfonate (PSS): names, main component and additives.

**Table S2.** Average number of hydrogen bonds (NHb) between peptides and the PSS_20 molecule, per single PSS group (MD results for 1:1 PSS-peptide systems at different salt concentrations). Results are averaged over the last 200 ns of simulations.

| NaCl concentration<br>mM | System | NHb |
| --- | --- | --- |
| 50 | 1 PSS_20 - 1 GR10 | $0.67 \pm 0.11$ |
| | 1 PSS_20 - 1 PR10 | $0.52 \pm 0.14$ |
| | 1 PSS_20 - 1 PA10 | $0.05 \pm 0.05$ |
| 150 | 1 PSS_20 - 1 GR10 | $0.58 \pm 0.11$ |
| | 1 PSS_20 - 1 PR10 | $0.45 \pm 0.12$ |
| | 1 PSS_20 - 1 PA10 | $0.04 \pm 0.04$ |
| 500 | 1 PSS_20 - 1 GR10 | $0.57 \pm 0.10$ |
| | 1 PSS_20 - 1 PR10 | $0.41 \pm 0.10$ |
| | 1 PSS_20 - 1 PA10 | $0.04 \pm 0.04$ |

**Table S3.** Average number of hydrogen bonds (NHb) between peptides and the PSS_20 molecules, per single PSS group (MD results for 7:7 PSS-peptide systems at different salt concentrations). Results are averaged over the last 50 ns of simulations.

| NaCl concentration<br>[mmol/l] | System | NHb |
| --- | --- | --- |
| 50 | 7 PSS_20 - 7 GR10 | $0.72 \pm 0.07$ |
| | 7 PSS_20 - 7 PR10 | $0.67 \pm 0.07$ |
| | 7 PSS_20 - 7 PA10 | $0.04 \pm 0.02$ |
| 150 | 7 PSS_20 - 7 GR10 | $0.71 \pm 0.07$ |
| | 7 PSS_20 - 7 PR10 | $0.65 \pm 0.08$ |
| | 7 PSS_20 - 7 PA10 | $0.04 \pm 0.02$ |
| 500 | 7 PSS_20 - 7 GR10 | $0.74 \pm 0.08$ |
| | 7 PSS_20 - 7 PR10 | $0.63 \pm 0.07$ |
| | 7 PSS_20 - 7 PA10 | $0.05 \pm 0.02$ |

